# Armed Macrophages as Hunters for Photodynamic Therapy of Systemic Bacterial Infections by Bathing in the Sunshine

**DOI:** 10.1101/2024.01.04.574019

**Authors:** Zehui Wang, Lai Wang, Lin Zhou, Xinfu Zhang, Yi Xiao

## Abstract

There has been a vision to conduct therapy using sunlight since ancient Greece, Egypt, and India. In modern medicine, photodynamic therapy is one popular form of therapy that uses light to excite photosensitizers to eliminate malignant and other diseased cells. It offers highly effective and broad-spectrum therapeutic efficacy. Yet, there are several hindrances to a real treatment of disease through photodynamic therapy, such as the limitation on the irradiation depth and areas, the regulation of side effects, etc. Usually, the patients should be kept in a dark environment during and after the therapeutic process for days to avoid side effects induced by light in daily life, not mention to conduct the phototherapy through sunbathing. Based on the above consideration, we propose an innovative idea to bring photodynamic therapy back to the origin of phototherapy-bathing in the sunshine. Namely, we designed a “live drug”, as a smart hunter, named A-RAWs, by loading an “off-on” type of photosensitizer in macrophages to solve two technical problems. Firstly, to find and capture bacteria accurately, and then transport these bacteria to the epidermis through the blood capillary, where sunlight is reachable. Secondly, to minimize the potential side effects and maximize the therapeutic time windows and efficacy by using bacteria as the trigger of the photodynamic effect. Specifically, we first designed and synthesized a lysosome-targeted and inflammation-activated NIR photosensitizer (Lyso710A), and loaded it in the lysosomes of macrophages. These “armed” macrophages were transferred into the infected host to capture deep-tissue bacteria through innate immunity and transport the captured bacteria through the bloodstream to superficial skin. Finally, the pathogenic bacteria are killed by the photodynamic effect when transported to the epidermis and receive sunbathing. In vivo experiment demonstrates a 100% therapeutic efficiency on systemic bacterial infection model. We also demonstrated the working mechanism of this “live drug” using a lung infection model. This hunter demonstrates high intelligence to break the limitations of current photodynamic therapy and executes photodynamic therapy of deep-tissue bacterial infection simply by bathing in the sunshine.

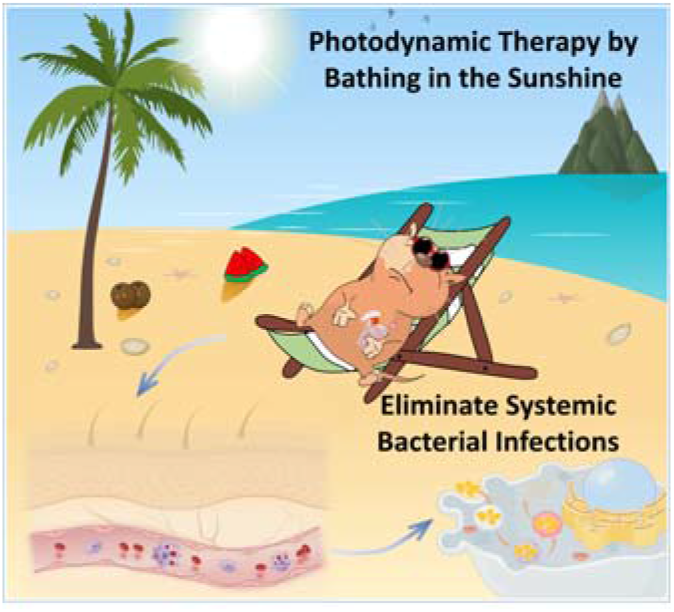

## Introduction

There has been a vision to conduct therapy using sunlight since the beginning of human civilization. The earliest record of phototherapy, known as heliotherapy, can be traced back to ancient Greece, Egypt, and India^1, 2^. People use sunlight as a therapeutic agent to treat various diseases, such as pain management, hair growth, skin treatments, etc^3–5^. In modern medicine, phototherapy is still present in treating skin diseases, such as leukoderma, tinea, and others^6–8^. Photodynamic therapy is one popular form of phototherapy that uses light to excite photosensitizers to generate reactive oxygen species (ROS), which are highly effective in destroying bio-macromolecules and eliminating malignant and other diseased cells. This approach offers highly effective and broad-spectrum therapeutic efficacy. However, the ROS may also damage healthy cells if not appropriately regulated. Generally, several limitations hinder the extensive clinical translation of photodynamic therapy. The first and most important one is that current phototherapy is limited to superficial areas due to the limitation of irradiation depth of light^9, 10^. Researchers have explored a variety of radiation strategies to promote the performance of photodynamic therapy in deep tissues, including implantable luminescence, chemiluminescence bioluminescence, etc^11–13^. Some modalities demonstrated improved therapeutic efficacy for deep lesions, but these techniques are somewhat complicated and less universal^14^. The second one is the challenges of delivering photosensitizers to diseased cells, or mobile pathogens, such as a bacterium, that moves freely in the body of a patient, as well as restricting the accumulation of photosensitizers in healthy cells^9, 15^. The methods of drug delivery such as topical sprays and injections have demonstrated good therapeutic efficacy in models of lesions in specific confined spaces, including wounds and tumors, but do not apply to real clinical conditions^16, 17^. The third is that most photosensitizers are of the “always on” type and the off-target photosensitizers retained in the skin and blood capillary can be excited by the daily light, which may result in damage to healthy tissue. Therefore, patients are forced to avoid prolonged exposure to sunlight even after the treatment to reduce potential side effects ^18–20^. These limitations make current photodynamic therapy far different from heliotherapy, and even can be regarded as anti-heliotherapy. Solutions to promote the targeting accuracy and the specificity (for example turn-on type) of photosensitizers to diseased cells are needed, to simplify and translate photodynamic therapy into a clinically executable treatment, preferably using sunlight as the light source.

We have been expecting a smart hunter who can carry photosensitizer, capture mobile pathogens deep in the body of patients, and transport these pathogens to the surfer layer of the body, where sunlight is reachable. To the best of our knowledge, there hasn’t been any report on such smart tools and photodynamic therapy for deep-tissue diseases using sunlight. However, firstly, from the aspect of targeting and transportation of pathogens, we do have such a smart tool of our own, that is the natural immune system. Macrophages can track, engulf, digest, and transport foreign substances. They recognize and capture pathogens, such as bacteria, by tracking cytokines secreted by them via the blood and body fluid system in extremely efficient and accurate patterns^9, 10, 21^. They migrate and transport nondegradable substances outside of the body via the blood and body fluid system^22^. Therefore, the chemotactic feature of macrophages endows the smartest and most accurate targeting and transportation ability as we expected. Secondly, restricting the damage by the photosensitizer on healthy tissue is as important as conversing their efficiency to efficacy. Our expectation is that the photodynamic effect of the photosensitizers stays “off” stability before recognizing pathogens. Light irradiation (including sunlight) will excite the photosensitizer that is turned on by the pathogens but not the photosensitizers in the off-state. In this circumstance, even sunlight (which may irradiate the whole body) can be used as a light source for photodynamic therapy. Such a turn-on mechanism can prolong the lifetime of macrophages loaded with photosensitizers by keeping them on standby.

To verify the idea of using macrophages as smart soldiers to carry photosensitizer, capture and transport mobile pathogens, and eliminate them through sunbathing, we built a septicemia mice model (systemic bacterial infection) and performed photodynamic therapy by simply bathing the infected mice in the sunshine. Septicemia is a life-threatening condition caused by whole-body bacterial infection, which leads to tissue damage, organ failure, and 30-50% death, particularly in people with chronic diseases or weakened immune systems^10, 23, 24^. Photodynamic therapy has been well established in numerous epidermal and limited area antibacterial infection models^25–31^. The challenges to cure septicemia include: firstly, it is a whole-body infection, that bacteria distribute in the bloodstream and all organs. Secondly, it progresses rapidly therefore leaving limited time for the identification of the bacterial species before the administration of the right antibiotic. Thirdly, severe side effects and the emergence of drug-resistant bacteria make treatments based on antibiotics fail sometimes. Given the above considerations, we designed a “live drug”, named A-RAWs, that is smart to find and capture bacteria autonomously, turn on its photodynamic effect upon capturing bacteria, and then cure systemic bacterial infection through sunbathing. Specifically, we first designed and synthesized a lysosome-targeted activatable NIR photosensitizer (Lyso710A). This photosensitizer was then loaded into the lysosome of macrophages through standard culturing. Then these “armed” macrophages were transferred into the infected host to capture deep-tissue bacteria through innate immunity and transport the captured bacteria through the bloodstream to superficial skin partially. Lyso710A stays in the off-state (weak photodynamic effect) in the lysosome for more than three days till it is turned on (strong photodynamic effect) upon the oxidative by the endogenous hypochlorite acid released by macrophages when capturing bacteria. Finally, the bacteria are killed by the photodynamic effect when they are transported to the epidermis and receive sunbathing. Importantly, the photodynamic effect is restricted within each macrophage, which avoids damaging healthy cells in the neighborhood. This “live drug” demonstrates high intelligence to break the limitations of current photodynamic therapy and executes photodynamic therapy of deep-tissue bacterial infection simply by bathing in the sunshine.

## Results and Discussion

### 1. The construction of A-RAWs: molecular design and cell engineering

We first designed and synthesized a lysosome-targeted near-infrared (NIR) photosensitizer (Lyso710A) that affords a stable lysosome targeting ability, a specific response toward hypochlorous acid and high photodynamic effect after turned on by hypochlorous acid. As shown in Figure 1a, two phenothiazines were conjugated to a BODIPY core through Knoevenagel condensation to extend its absorption wavelength to the NIR region and quench the photodynamic effect through Photoinduced Electron Transfer (PET) effect, two iodine atoms were introduced at the 2,6-position of the BODIPY core to maximize the photodynamic effect, one tertiary amine moiety was grafted onto each phenothiazine to afford lysosome targeting ability and a short PEG chain was linked to improving its bio-compatibility. In this design, the oxidation of the thioether (by the endogenous hypochlorous acid within the macrophage) will restrict the PET effect therefore resulting in the recovery of its fluorescence and photodynamic effect (Fig. 1a). According to the physiological feature of macrophages, the intracellular oxidative chain burst (e.g., elevated hypochlorous acid levels) is a typical response toward bacterial infection. Therefore, these ROS, especially the hypochlorous acid, can work as the trigger of the photodynamic effect of Lyso710A. Further on, the stable lysosomal targeting ability, high photodynamic efficiency and strong NIR absorption are essential properties to achieve precise and efficient antimicrobial activity (Fig. 1b). The ^1^O_2_ is a highly reactive/destructive species but with a short half-life period (∼0.6 µs) and diffusion radius (∼20 nm)^32, 33^. Two tertiary amine groups enable the lysosome targeting, which may pull the photosensitizer and bacteria within the same lysosome therefore fully transferring its reactivity to therapeutic efficacy. Additionally, its near-infrared absorption is in favor of phototherapy at a deeper level reaching the blood capillary. In general, Lyso710A shows three features: i) specific hypochlorous acid response, ii) stable lysosomal localization, and iii) high photodynamic efficiency post-activation. The structure of Lyso710A has been characterized by ^1^H NMR and HRMS (SI Fig. 12-15).

**Fig. 1.**
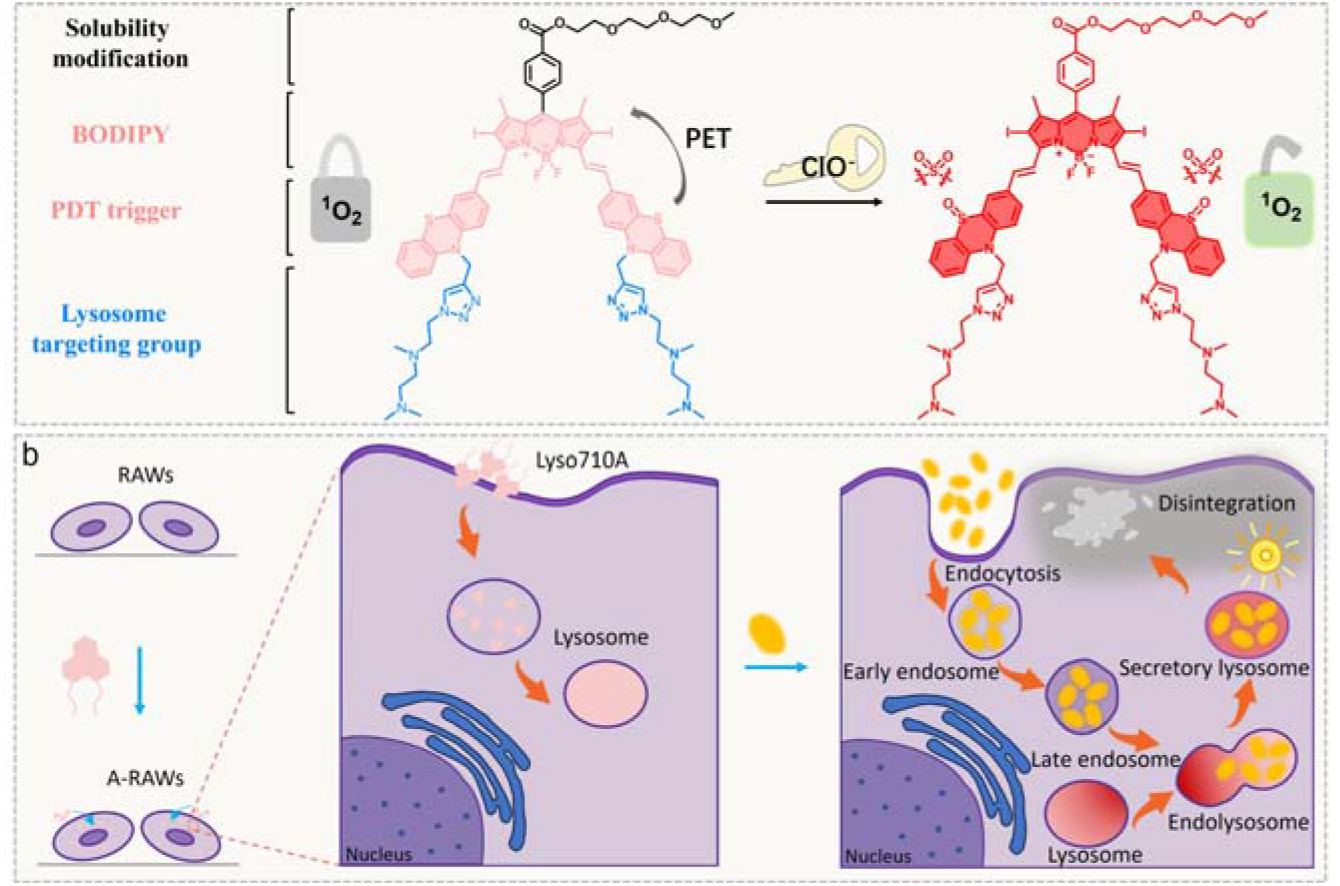
(a) The structure and working mechanism of Lyso710A. Oxidative of the phenothiazines turns on the photodynamic effect by breaking the photoinduced electron transfer process. (b) The construction procedure and working mechanism of A-RAWs.

We second fabricate the final live drug simply by loading Lyso710A into macrophages through culturing at 5 µM for 3 h. These armed macrophages (A-RAWs) stay in “stand-by” mode, showing no photodynamic effect (Fig. 1b). They are ready to be administrated into the infected host and remain alive in “stand-by” before the uptake of bacteria turns on the photodynamic effect of Lyso710A. The synthetic route and preparation method of A-RAWs is illustrated in the supporting information (SI Fig. 1).

### 2. Optical Properties and Photoactive Performance

Next, we evaluate the basic photophysical properties of Lyso710A to determine its suitability for antimicrobial applications. As shown in Figure 2a, Lyso710A exhibits intense absorption in the entire spectral region ranging from 350 to 800 nm with a maximum extinction coefficient of 84450 M^−1^·cm^−1^ at 713 nm in PBS (with 0.1% Triton X-100). This broad absorption spectrum indicates the advantage of Lyso710A to absorb sunlight for photodynamic therapy. Lyso710A shows no detectable emission due to the PET effect from the phenothiazines. Then, we examined the selectivity and responding rate of the Lyso710A for hypochlorous acid (HClO). Upon adding HClO into the aqueous solution of Lyso710A (10 μM), the maximal absorption band shifts from 713 nm to 691 nm, and an emission peak at 715 nm appears (Fig. 2a and 2b). The recovery of the fluorescence is positively correlated with the amount of hypochlorous acid, demonstrating a fluorescence quantum yield of 0.4% (Fig. 2b inset). The kinetic curve of fluorescence intensity (at 715 nm) upon the addition of 100 µM of HClO shows that the oxidation can be completed within 8 s (Fig. 2c). To evaluate the selectivity of Lyso710A toward HClO, we added excessive other reactive oxygen and nitrogen species (ROS/RNS, 1.5 mM), into the aqueous solution of Lyso710A (10 μM). As shown in SI figures 3a and 3b, the emission at 715 nm increases marginally when these ROSs and RNSs are added. This result shows that Lyso710A is highly selective for HClO, which guarantees a stable stand-by state and minimal photodynamic effect in macrophages before encountering bacteria. These photophysical properties show the specificity and response rate of Lyso710A toward HClO, which functions as an effective trigger of the photodynamic effect.

**Fig. 2.**
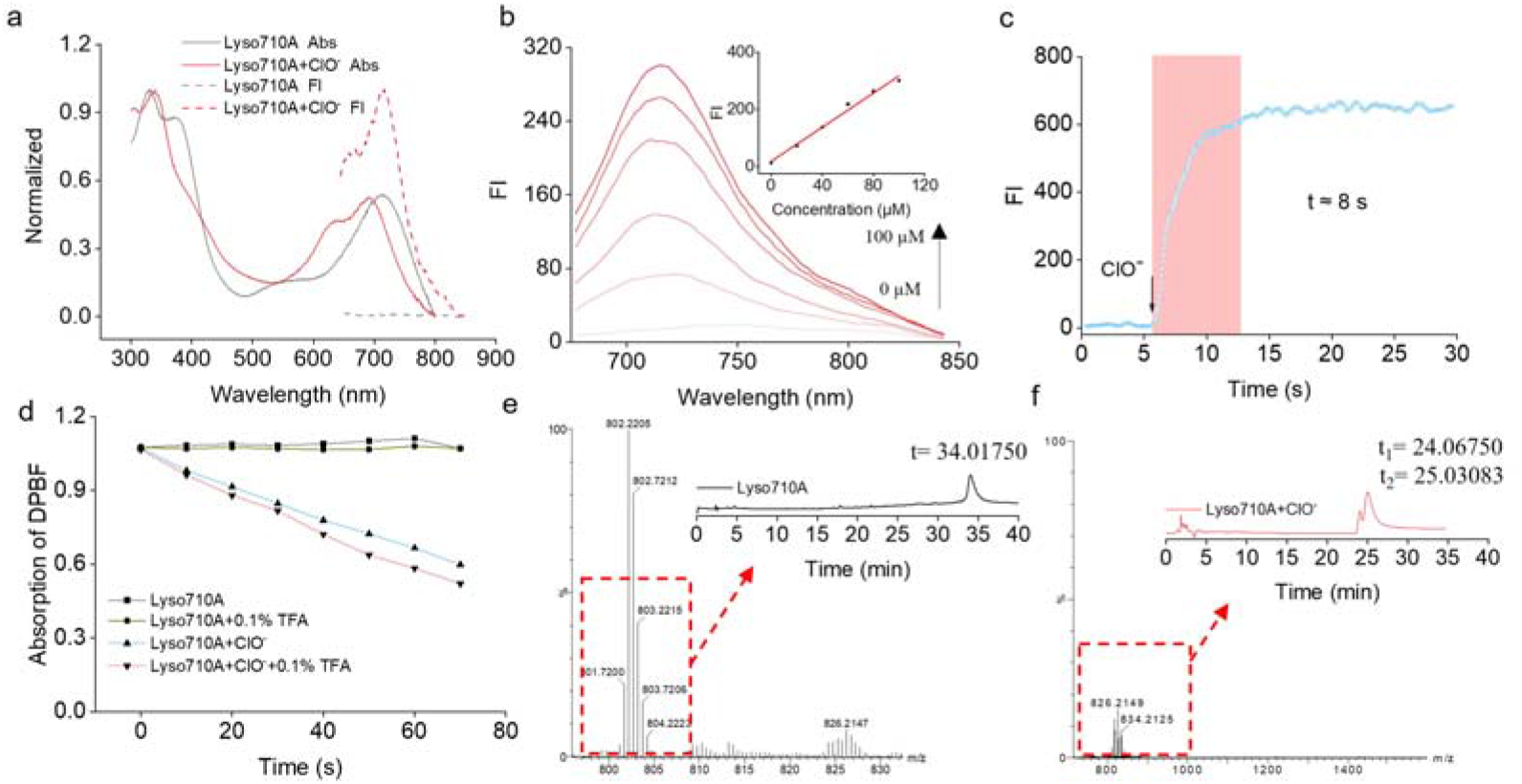
(a) The normalized absorption and emission spectrums of Lyso710A before and after adding HClO in PBS (0.1% Triton X-100). (b) Fluorescence spectra of Lyso710A (10 μM) upon the addition of HClO (0-100 μM) in PBS (0.1% Triton X-100). (c) Time-dependent plot: the fluorescent intensity of Lyso710A (10 μM) at 715 nm against time after the addition of HClO (100 µM) in PBS (0.1% Triton X-100). (d) Absorption intensity of DPBF at 415 nm against irradiation time (660 nm LED light) in the presence of oxidation products of Lyso710A induced by HClO in PBS (0.1% Triton X-100). (e) HRMS and HPLC analysis of Lyso710A. (f) HRMS and HPLC analysis of Lyso710A in the presence of HClO.

**Fig. 3.**
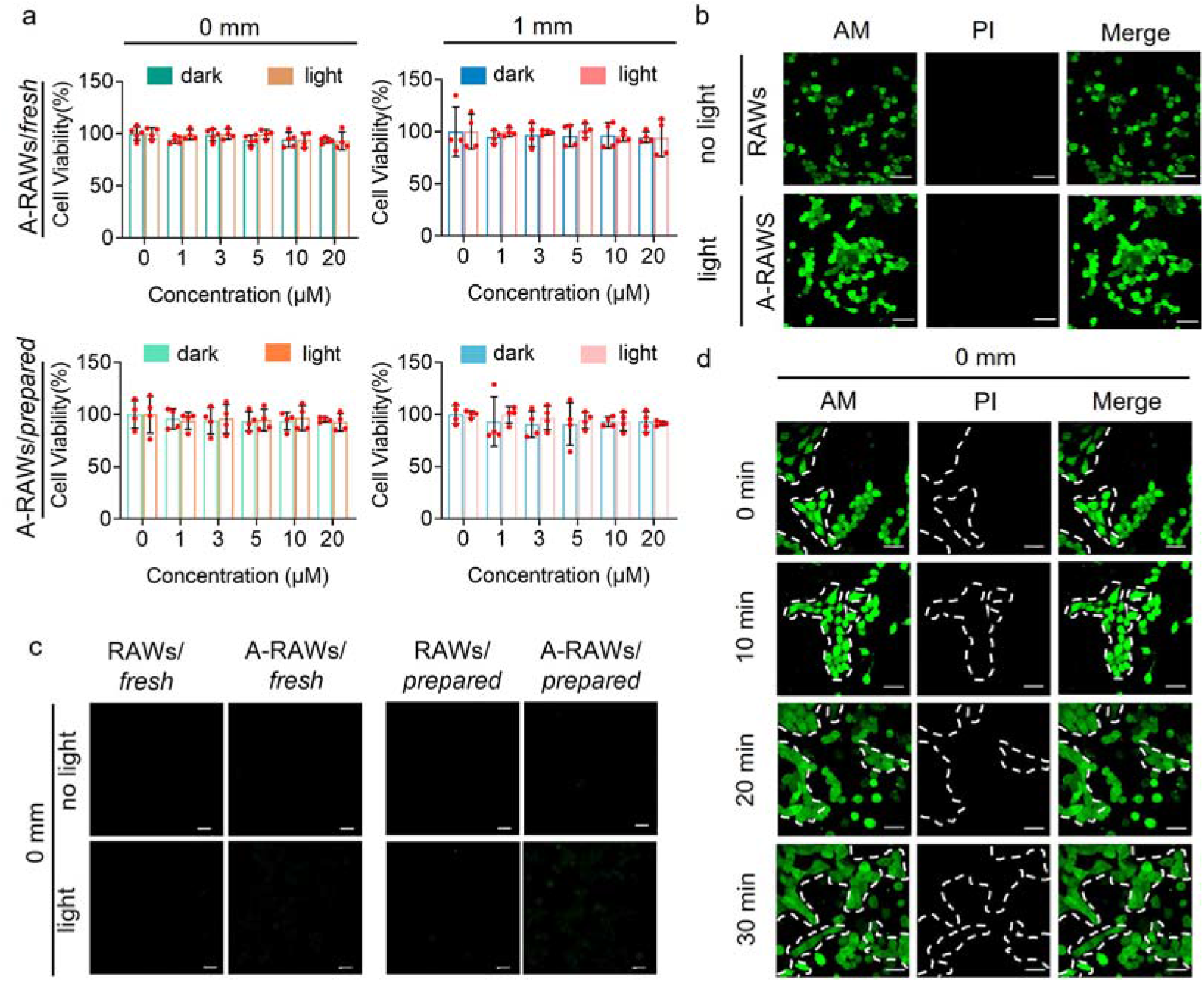
(a) A-RAWs/*fresh* and A-RAWs/*prepared* were evaluated for MTT survival at 0 mm and 1 mm thickness, respectively (dark or simulated sunlight t, 70 mW/cm^2^, 10 min). (b) Calcein-AM/PI fluorescence imaging assesses the cell viability before activation of A-RAWs/*fresh* with and without simulated sunlight (70 mW /cm^2^, 10 min). Bar: 30 µm. (c) Fluorescence imaging of intracellular ROS production monitored by DCF-DA after light/dark incubation with A-RAWs/*fresh* and A-RAWs/*prepared* through 0 mm thickness of chicken meat, respectively. Bar: 20 µm. (d) Calcein-AM/PI fluorescence imaging assesses the cell viability of co-cultured inactivated A-RAWs and COS-7 cells under simulated sunlight (70 mW/cm^2^, 10, 20, 30 min) through 0 mm thickness of chicken meat, respectively. The viability of cells was indicated through a Calcein-AM/PI co-staining assay. Bar: 30 µm.

### 3. Activation mechanism and photodynamic efficiency of Lyso710A

We then evaluated the activation mechanism and photodynamic efficiency of Lyso710A. According to the structural design, two conjugated phenothiazines quench the photodynamic effect through the PET effect, due to the electronic donating properties of the thioether. The oxidation of the thioether by hypochlorous acid will restrict the PET effect therefore resulting in the recovery of its fluorescence and photodynamic effect^34–36^. To clarify this mechanism, we first simulated the oxidation of Lyso710A using excessive amounts of HClO in solution and characterized the oxidation product through liquid chromatography and high-resolution mass spectrometry (HPLC and HRMS). As shown in Figure 2e and 2f illustration, the HPLC retention time for Lyso710A is 34.02 min, while the time for the oxidation products is 24.07 min and 25.03 min. The products from HPLC were used for mass spectra. As shown in Figure 2e, f, the m/z for Lyso710A ([M+2H]^2+^) is 802.2205, while the m/z for the oxidation products ([M+2H]^2+^) are 826.2149 and 834.2125 respectively, corresponding to the single oxidation of both phenothiazine groups (yielding sulfoxide) and double oxidation of both phenothiazine groups (yielding sulphone) HPLC and HRMS results indicate the successful oxidation of thioether into sulfoxide and sulfone. We further simulated the activation (oxidation) of Lyso710A in live RAWs with the bacterium (MRSA) and collected the oxidation products from cell lysate (see the method in SI) for characterization. The HRMS showed m/z ([M+2H]^2+^) of 810.2195 (single sulfoxide), 818.2175 (double sulfoxide), 826.2147 (single sulphone), and 834.2127 (double sulphone), corresponding to the four oxidation levels of the phenothiazine groups in Lyso710A (SI Fig. 4). This result indicates that the oxidative stress during bacterial infestation leads to the oxidation of Lyso710A into sulfoxide or/and sulfone forms.

Furthermore, we evaluated the photodynamic efficiency of Lyso710A before and after oxidation using DPBF as a ^1^O_2_ indicator. In neutral or 0.1% TFA/PBS, Lyso710A does not produce ^1^O_2_ (with an undetectable ^1^O_2_ quantum yield, Fig. 2d). While the ^1^O_2_ quantum yield of Lyso710A after oxidation was calculated to be 0.42 in neutral PBS (Fig. 2d). In acidic PBS, the number rise to 0.50 (Fig. 2d), which indicated tertiary amines quench the photodynamic effect to some extent in neutral conditions due to the extra PET effect, while releasing the photodynamic effect due to the ionization of the tertiary amines. This pH response is in favor of maximizing the photodynamic effect of the products of Lyso710A in the lysosome (pH∼4.0).

### 4. Durability and photo-toxicity of A-RAWs in standby mode

The turn-on mechanism of Lyso710A can prolong the lifetime of A-RAWs by keeping them in standby mode until they capture bacteria, which is one crucial property of performing phototherapy using weaker power but a longer period of light irradiation (such as sunbathing) and in favor of simplifying the therapeutic procedure, such as multiple phototherapies upon one administration of A-RAWs. According to the in vitro result, Lyso710A stays in the off state before oxidation, which means it shows no photo-toxicity with or without light irradiation. Therefore, A-RAWs may remain effective (in standby mode) in the host for days which is a long window period to capture most bacteria and executive phototherapy. In general, staying stably in standby mode before capturing bacteria is essential for A-RAWs to conduct phototherapy utilizing sunlight with weaker power but a longer irradiation period and to restrict damage to healthy cells in the neighborhood.

Given the above consideration, we evaluated the stability and the photo-toxicity of A-RAWs in standby mode (before capturing bacteria) to healthy cells in the neighborhood carefully. We first confirmed the cell viability of A-RAWs with and without light irradiation by MTT assay and live/dead cell co-staining assay. For the MTT experiment, as shown in Figure 3a, freshly prepared A-RAWs (named A-RAWs/*fresh*) show viability higher than 93.43% after being treated with Lyso710A in a concentration of up to 20 μM. The stained A-RAWs show a similar viability higher than 93.06% after light irradiation, indicating that A-RAWs are stable against light irradiation. We performed the same MTT assay under 1 mm of meat coverage to simulate the epidermis that covers blood capillaries. The viability of A-RAWs is up to 94.73% and 94.00% for groups with and without light irradiation. We performed the same assay using A-RAWs 72 h after the preparation (named A-RAWs/*prepared*), these A-RAWs/*prepared* (with or without meat coverage) show viability higher than 95.00% (20 μM group) without light irradiation. Light irradiation shows a viability (with or without meat coverage) higher than 92.61% (20 μM group). For live/dead cell co-staining assay, after light irradiation (simulated sunlight, 70 mW/cm^2^, 10 min), A-RAWs/*fresh* in standby mode shows intense fluorescence in Calcein-AM channel but no fluorescence in PI channel (Figure 3b), which indicates that Lyso710A in the off state shows undetectable toxicity to cells and A-RAWs can maintain alive and stay in standby mode under sunlight exposure.

To further demonstrate the durability (in view of photodynamic effect) of A-RAWs, we used DCFH-DA to detect ROS generation in freshly prepared A-RAWs (A-RAWs/*fresh*) and A-RAWs 72 h after preparation (A-RAWs/*prepared*). We first irradiated the RAWs and A-RAWs with simulated sunlight (70 mW/cm^2^, 10 min) and then performed fluorescence imaging to evaluate the fluorescence intensity of DCFH-DA. As shown in Figure 3c, no fluorescence was observed in the A-RAWs/*fresh* group, and A-RAWs/*prepared* group as in the RAWs of the control group, indicating that Lyso710A shows no photodynamic effect and stays in the off-state in A-RAWs for at least 72 h. We also performed the same assay through 1 mm of meat coverage. The same result was obtained that Lyso710A shows no photodynamic effect and stays in the off-state in A-RAWs for at least 72 h (SI Fig 5). These results confirm that A-RAWs are durable and maintained in standby mode for at least 72 hours, which provides a long therapeutic window period. To verify the photo-toxicity of A-RAWs to healthy cells in the neighborhood, we performed a co-culture assay using A-RAWs and COS-7 cells, which can be distinguished simply by their morphological features. 24 h after seeded, the co-cultured A-RAWs and COS-7 cells were irradiated using simulated sunlight at various times (70 mW/cm^2^, 10, 20, 30 min). Then, the viability of A-RAWs and COS-7 cells was monitored using a live/dead cell co-staining assay. Both A-RAWs (round cells) and adjacent COS-7 cells (spindle-shaped cells, encircled by dashed lines) showed strong green fluorescence in the Calcein-AM channel but no fluorescence in the PI channel upon 0-30 minutes of irradiation (Fig. 3d), which indicates both RAWs and COS-7 were alive. The same result was obtained when the irradiation was conducted through a 1 mm cover of chicken meat (SI Fig. 6). These results further indicate that A-RAWs are safe for healthy cells in the neighborhood and can remain alive and stay in a standby mode before being activated by bacteria.

### 5. Targeting stability and Photodynamic efficiency of A-RAWs in on-mode

After confirming the durability of A-RAWs, we investigated their targeting ability, response performance, and photodynamic efficiency towards bacteria. As the leading role of innate immune, macrophages play a critical part in nonspecific defense. They actively track and engulf bacteria into endosomes that fusion with lysosomes consequently^37^. Upon the phagocytosis of bacteria, macrophages are activated and promote oxidative stress by secreting hypochlorous acid, hydrogen peroxide, and other ROS^37–40^. Our design takes advantage of this immune response, specifically, by utilizing the ROS as a trigger to turn the photodynamic effect of Lyso710A on therefore turning A-RAWs into on-state. A-RAWs stay in a standby state until they capture bacteria, after which Lyso710A is gradually turned on (the oxidation of the phenothiazine part) in situ as intracellular oxidative stress is promoted. Finally, the Lyso710A in on-state eliminated the bacteria by generating ROS under light irradiation (Fig. 4a). The following experiments were performed to validate the above rationale. For all in vitro phototherapy assays, we added an extra group with a coverage of 1 mm chicken meet to simulate the sunlight irradiation of the blood capillary through the skin.

**Fig. 4.**
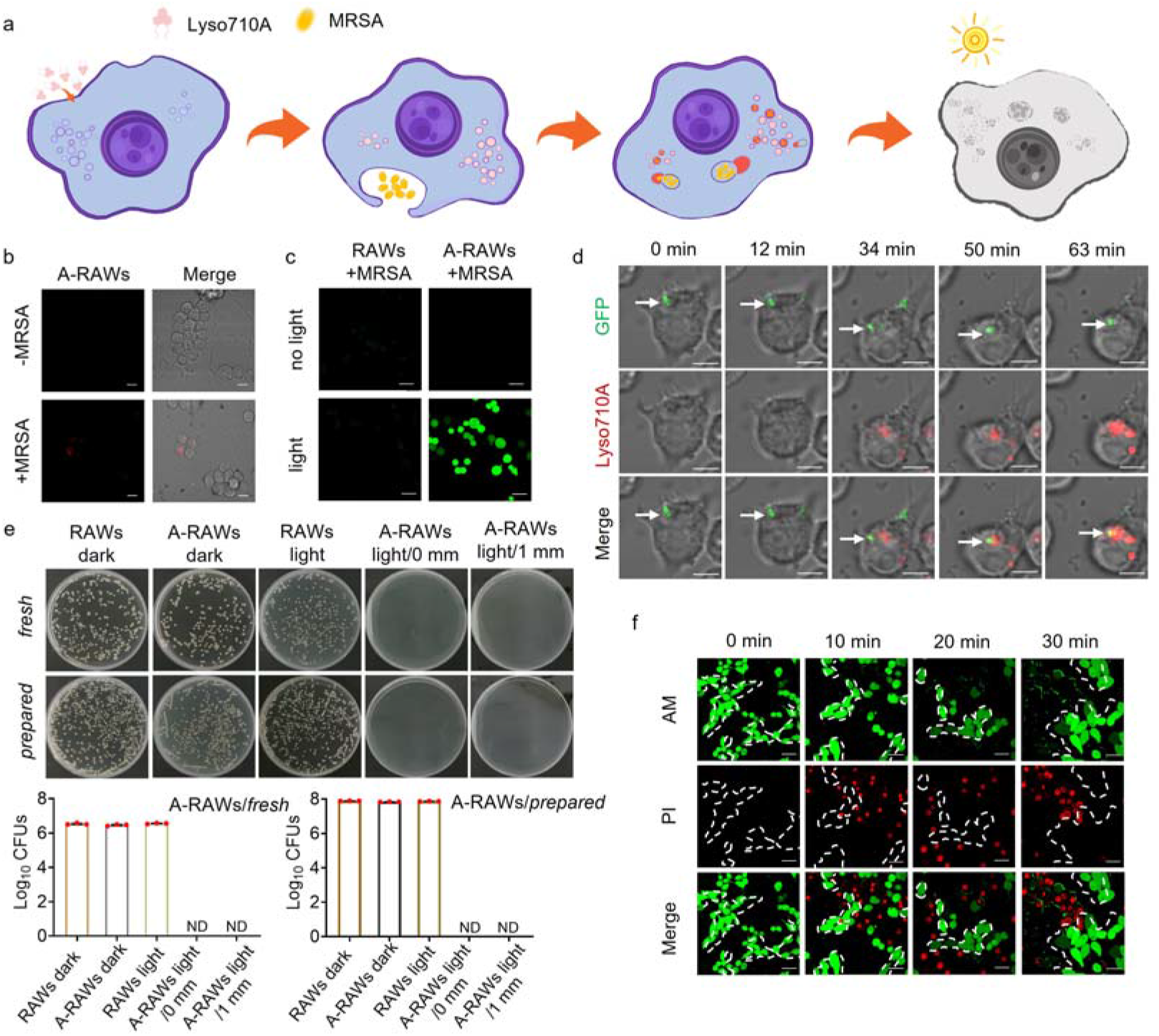
(a) Schematic diagram of A-RAWs capturing MRSA and activating Lyso710A. (b) Fluorescence imaging of A-RAWs before and after MRSA stimulation. Bar: 10 µm. (c) Fluorescence imaging of intracellular ROS production monitored by DCFDA after no/light incubation with A-RAWs after MRSA stimulation (dark or simulated sunlight t, 70 mW/cm^2^, 10 min). Bar: 10 µm. (d) Confocal images of A-RAWs capture MRSA and activate Lyso710A. Bar: 10 µm. (e) Plate count evaluates the survival of MRSA in intracellular post-phototherapy using A-RAWs/*fresh* and A-RAWs/*prepared* with different thickness coverage (70 mW/cm^2^, 40min). (f) AM/PI fluorescence imaging assesses the cell viability of co-cultured activated A-RAWs and COS-7 cells under simulated sunlight (70 mW/cm^2^, 10, 20, 30 min). Bar: 30 µm.

Firstly, we evaluated the targeting ability of Lyso710A toward lysosomes and demonstrated the response of A-RAWs toward bacteria. A-RAWs were treated with HClO carefully to turn the fluorescence of Lyso710A on and then used for co-localization imaging with LysoTracker®Green DND-26. As shown in SI Figure 7, co-localization imaging shows a good overlay between the fluorescence from LysoTracker®Green DND-26 and oxidized Lyso710A with a Pearson Correlation Coefficient of 0.95, indicating the targeting specificity of Lyso710A toward lysosomes. This lysosomal targeting specificity is another crucial property to apply the reactive ^1^O_2_ on bacteria therefore conducting precise and efficient photodynamic therapy.

Further on, we evaluated the response of A-RAWs toward bacteria by comparing the fluorescence intensity and the ability to generate ROS before and after the capture bacteria. As shown in Figure 4b, A-RAWs show no fluorescence before being treated with bacteria, but intense red fluorescence after they capture bacteria, which confirms that A-RAWs turn into on-state when they capture bacteria. Simultaneously, the generation of ROS within A-RAWs was monitored using a commercial indicator (2’, 7’-dichlorodihydrofluorescein diacetates, DCFH-DA). As shown in Figure 3d, after being treated with MRSA, no green fluorescence of DCF-DA is observed in the control group (RAWs without Lyso710A) with or without light irradiation (simulated sunlight, 70 mW/cm^2^, 10 min), indicating no ROS is generated in these groups. However, the strong green fluorescence of DCF-DA is observed in A-RAWs after they are treated with MRSA and then irradiated with light (Fig. 4c), indicating Lyso710A is turned on and generates ROS upon light irradiation. This result suggests that A-RAWs are in on-mode and show enhanced photodynamic effects after they capture bacteria. We further imaged the phagocytosis process of MRSA by A-RAWs to demonstrate its response to bacteria. As shown in Figure 4d, MRSA (pseudo color green, stained by membrane tracker Mem-SQAC) is taken into the cytosol by A-RAWs started at the 12^th^ minute. As the endosomes mature, the fluorescence from the oxidized Lyso710A (red) increases progressively at the 34^th^ minute. Finally, the MRSA is transported to the lysosome at the 63^th^ minute, as indicated by the overlay of the green fluorescence of bacteria and the red fluorescence from the oxidized Lyso710A. This dynamic visualization confirms the feasibility of our strategy and the response of A-RAWs towards bacteria.

Thirdly, we tested the antimicrobial efficiency of A-RAWs (in on-mode) in vitro. We chose MRSA as a representative bacterium and treated A-RAWs with MRSA for 0.5 h to enable the phagocytosis of bacteria. We then removed the free bacteria in the culture medium using gentamicin sulfate (100 g/mL) for 1 h. We next treated these A-RAWs that had taken bacteria with simulated sunlight (70 mW/cm^2^, 40 min). These A-RAWs were lysed after phototherapy and used for culturing bacteria colonies. According to the colony counts (Fig. 4e), MRSA in the test group (A-RAWs light/0 mm) was eliminated. To simulate the epidermis that covers blood capillary, we performed the same antimicrobial assay through 1 mm chicken meat. As shown in Figure 4e, MRSA is also completely removed (A-RAWs light/1 mm). By contrast, RAWs groups (dark and light) showed a CFU of at least 3.43×10^6^. The A-RAWs dark group showed a CFU of 2.91×10^6^, indicating the efficacy of phototherapy. To demonstrate the durability and antimicrobial window period of A-RAWs, we used A-RAWs/*prepared* to perform the same in vitro assay. MRSA in two tested groups (A-RAWs light/0 mm and A-RAWs light/1 mm) were also eliminated. These results demonstrate that A-RAWs have a long therapeutic window of up to 72 h and a high therapeutic efficiency of over 1 mm of coverage.

Lastly, safety is a critical precondition for photodynamic therapy using sunlight. Photodynamic therapy offers high therapeutic efficiency, while the ROS may also damage healthy cells if not appropriately regulated. In other words, minimizing photodamage to healthy tissue is as important as maximizing photodynamic efficiency in the focus. We therefore evaluate the phototoxicity of A-RAWs in on-state to healthy cells in the neighborhood through a co-culture assay. We co-cultured A-RAWs (pre-activated using a low concentration of HClO) and COS-7 cells to simulate the in vivo situation that A-RAWs are surrounded by healthy cells. We then irradiated these co-cultured cells with simulated sunlight (70 mW/cm^2^) for 10 min, 20 min, or 30 min, and assessed the viability of cells using live/dead cell co-staining assay. As shown in Figure 4f, A-RAWs in on-mode (round cells) showed progressive apoptosis as indicated by decreased green fluorescence in the Calcein-AM channel and increased red fluorescence in the PI channel. But the adjacent COS-7 cells (spindle-shaped cells, encircled by dashed lines) show strong green fluorescence in the Calcein-AM channel and no fluorescence in the PI channel upon 30 minutes of irradiation (Fig. 4f), which means these COS-7 cells are still alive. This result indicates an efficient photodynamic effect of A-RAWs in on-mode, but the ROS generated by the on-state Lyso710A are restricted within A-RAWs and are harmless to healthy cells in the neighborhood.

### 6. Treatment of systemic bacterial infections in Septicemia mice model through sunbathing

Given the promising in vitro result, we finally utilized A-RAWs for the treatment of septicemia through sunbathing. Septicemia is a life-threatening condition that arises from a serious whole-body infection. It requires immediate treatment with intravenous fluids and a large dose of broad-spectrum antibiotics. However, according to an updated report, the risk of death from septicemia is as high as 30%, while for severe septicemia it is as high as 50%^41^. Our proposal for the “live drug”, A-RAWs, is designed to supplement the suppressed innate immunity, and capture and eliminate bacteria through photodynamic effect. A-RAWs maintain the feature of macrophages that can track and capture bacteria and lead to accurate photodynamic therapy within single RAWs. Therefore, phototherapy can be conducted using a very small dose of photosensitizer^9^. More importantly, the “off-on” trigger of the photodynamic effect of Lyso710A makes our therapy highly bacteria-specific, and safe for healthy cells, therefore can be performed through sunbathing. To the best of our knowledge, this is the first report of photodynamic therapy by bathing the whole body in the sunshine. To assess the therapeutic efficacy of A-RAWs on MRSA-induced septicemia, a “sunbathing” therapy was demonstrated.

We first induced immunodeficiency in mice by administering cyclophosphamide (CY) for three consecutive days, which was confirmed by the low levels of white blood cells (WBCs) and lymphocytes (LYMs) (Fig. 5c, d). We then established the systematic infection by injecting MRSA intravenously (Fig. 5a, method see SI). 2 hours after injecting MRSA (early systemic infection), the infected mice were divided into four groups, namely, the PBS, RAWs, Lyso710A, and A-RAWs groups and intravenously injected with PBS, RAWs, Lyso710A, and A-RAWs correspondingly. The equivalent dose of Lyso710A in the Lyso710A group and the A-RAWs group are both 0.14 mg/Kg mice (For detailed calculations refer to the supporting information, SI Fig. 2). We then placed these mice in sunlight at daytime (6 hours/day, detected power density: 50-120 mW/cm^2^) for three days to conduct phototherapy (a standard method of irradiance in this study, method see SI). The mice used to monitor bacterial load before and after treatment are described as follows: for the A-RAWs group, mice were killed immediately after treatment. For the PBS, RAWs, and Lyso710A groups, samples were taken immediately at the time of death, as they died during the sunlight period. All mice samples were used for evaluating the bacteria load through colony counting (method see SI). We found that the bacteria in the blood system of the A-RAWs group were eliminated, through sunbathing, and curing the infected mice in the end (Fig.5b). Only a small number of bacteria (less than 0.097×10^5^ CFUs) can be found in the liver, kidney, or spleen, that was completely removed in 20 days of recovery (SI Fig.8). By contrast, in the PBS group, rapid loss in body weight (ΔBWs over 3 g in 48 h) and quick death in 100% were observed within 48 hours. MRSA had proliferated rapidly in the blood (2.14×10^5^ CFUs) and main organs of mice (Heart: 4.28×10^5^ CFUs; Liver: 9.36×10^5^ CFUs; Spleen: 5.38×10^5^ CFUs; Lung: 2.97×10^5^ CFUs; Kidney: 4.58×10^5^ CFUs, Fig. 5b, e) when they died. Mice in the RAWs group, showed slightly extended survival time upto 60 hours, but still developed into a systemic bacterial infection and died with high bacteria load (Blood: 1.70×10^5^ CFUs; Heart: 5.38×10^5^ CFUs; Liver: 4.36×10^5^ CFUs; Spleen: 4.42×10^5^ CFUs; Lung: 1.26× 10^5^ CFUs; Kidney: 1.96×10^5^ CFUs, Figure 5b, e), which indicates the limited defense efficiency of natural immune cells toward a severe bacterial infection. For mice in the Lyso710A group, the phototherapy failed to inhibit MRSA proliferation either in the blood (1.20×10^5^ CFUs) or in main organs (Heart: 5.29×10^5^ CFUs; Liver: 2.77×10^5^ CFUs; Spleen: 2.78×10^5^ CFUs Lung: 0.85×10^5^ CFUs; Kidney: 5.06×10^5^ CFUs), resulting in 100% of death within 48 hours (Fig. 5b, e, SI Fig. 8). Although the equivalent dose of Lyso710A in the Lyso710A group and the A-RAWs group. at the same, the free Lyso710A showed no efficacy at all, which highlights the necessity of targeting accuracy for efficient therapy. These results prove the ide a of a “live drug” and highlight the successful application of A-RAWs in photodynamic therapy of systematic infection through sunbathing. Finally, in an extra A-RAWs group for recording the survival rate, mice were cured at 100% 30 days after the treatment. These mice showed a complete return to a healthy state, including recovery of body weights (BWs>23 g), white blood cells (WBCs=6.34×10^6^ ml^-1^), and lymphocytes (LYMs=5.10×10^6^ ml^-1^) (Fig. 5 c, d, e, f). H&E staining confirmed the restoration of pathological organs in “restored” immunocompetent mice (SI Fig. 9).

**Fig. 5.**
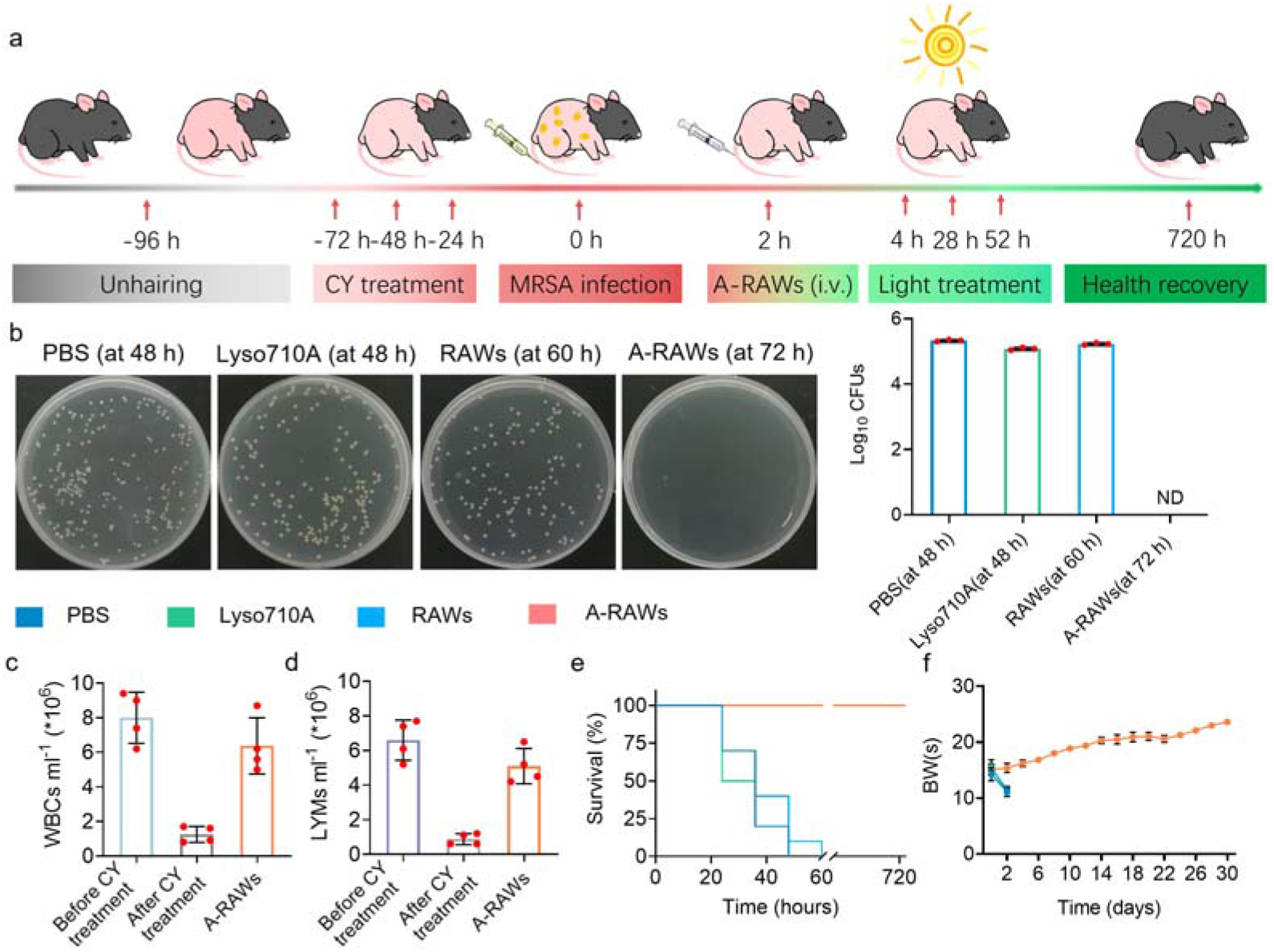
(a) Schematic diagram of the treatment time for systemic bacterial infection caused by MRSA mixture injection in C57BL/6 mice. (b) Plate counting of bacterial load in the blood at sunlight treatment three days (6 hours/day). (c-d) WBCs (c) and LYMs (d) of mice. (e) The survival rate of mice with epidermal bacterial infection post phototherapy. (f) BWs of mice.

### 7. Transportation of MRSA from the infection sites to the blood capillary in the epidermal layer

It is a universal limitation of phototherapy, that the penetration depth of light is only several millimeters. It decides that photodynamic therapy is currently only applicable to superficial focus. In the septicemia mice model in the last section, bacterial infections rapidly spread from the bloodstream to tissues and organs as indicated by the colony counts (SI Fig. 10). However, post the administration of A-RAWs, sunbathing on the skin of mice was eliminated bacteria not only in the blood system but also in the organs deep in the body. According to the literature, immune cells have the ability to travel and migrate throughout the organs and tissues^42^. They can capture and transport substances back into the bloodstream and body fluid circulation. For example, Cook et al. studied the trafficking of dendritic cells in the lung and found that after capturing antigens, these cells migrated through lymphatic vessels to nearby lymph nodes and interacted with other immune cells to initiate immune responses^43^. Similarly, Randolph et al. found that macrophages captured and transported lipids through lymphatic vessels to draining lymph nodes to the spread of inflammation and the progression of atherosclerosis^44^. Based on these former researches, we hypothesize that macrophages may track and capture bacteria in deep tissue, then transport the captured bacteria to other sites via the circulatory system, for example through blood capillary. The transportation of the captured bacteria in the blood capillary in the epidermal layer provides a perfect therapeutic time window and a large irradiation area for phototherapy through sunbathing.

To explore the mechanism and demonstrate the rationale of our therapeutic strategy, we built a lung infection mice model as a limited infection site deep in the body and administrated A-RAWs into these mice. We then monitored the distribution pattern of A-RAWs as well as the bacteria, focusing on identifying A-RAWs with bacteria intracellular (capturing bacteria) in the blood. If we identify A-RAWs with bacteria intracellular in the blood but no free bacteria in the blood, we may decide that A-RAWs capture bacteria in the lung and transport the bacteria into the blood system. According to a former study, bacteria-induced pneumonia in immunocompromised hosts may progress to systemic infection (bacteria spread through the body) 24 h after the infection^45^. We first confirmed this proliferation feature of bacteria from the lung to the whole body. We collected samples of heart, liver, spleen, lung, kidney and blood for bacterial enumeration at 4 h and 8 h after nasal administration of MRSA to immunodeficient mice (Fig. 6b, SI Fig. 11). According to plate counting, we found plenty of bacteria in the lungs with a CFU of 0.98×10^5^, but no bacteria in the blood and other major organs at 4 h after the infection, which indicated the lung infection had not spread in 4 h (Fig. 6b, SI Fig. 11). At 8 h after the infection, we found the bacteria in the lung were as high as 4.63×10^5^ CFU, but still no bacteria in other major organs. However, we found a trace number of bacteria in the blood with a low CFU of 200. This small number of bacteria in the blood indicated the lung infection had spread to the blood at that time. Therefore, we decided to monitor the presence of A-RAWs with bacteria intracellular in the blood within 4 h since the infection, in order to avoid the situation that A-RAWs capture bacteria that has already spread into the bloodstream.

**Fig. 6.**
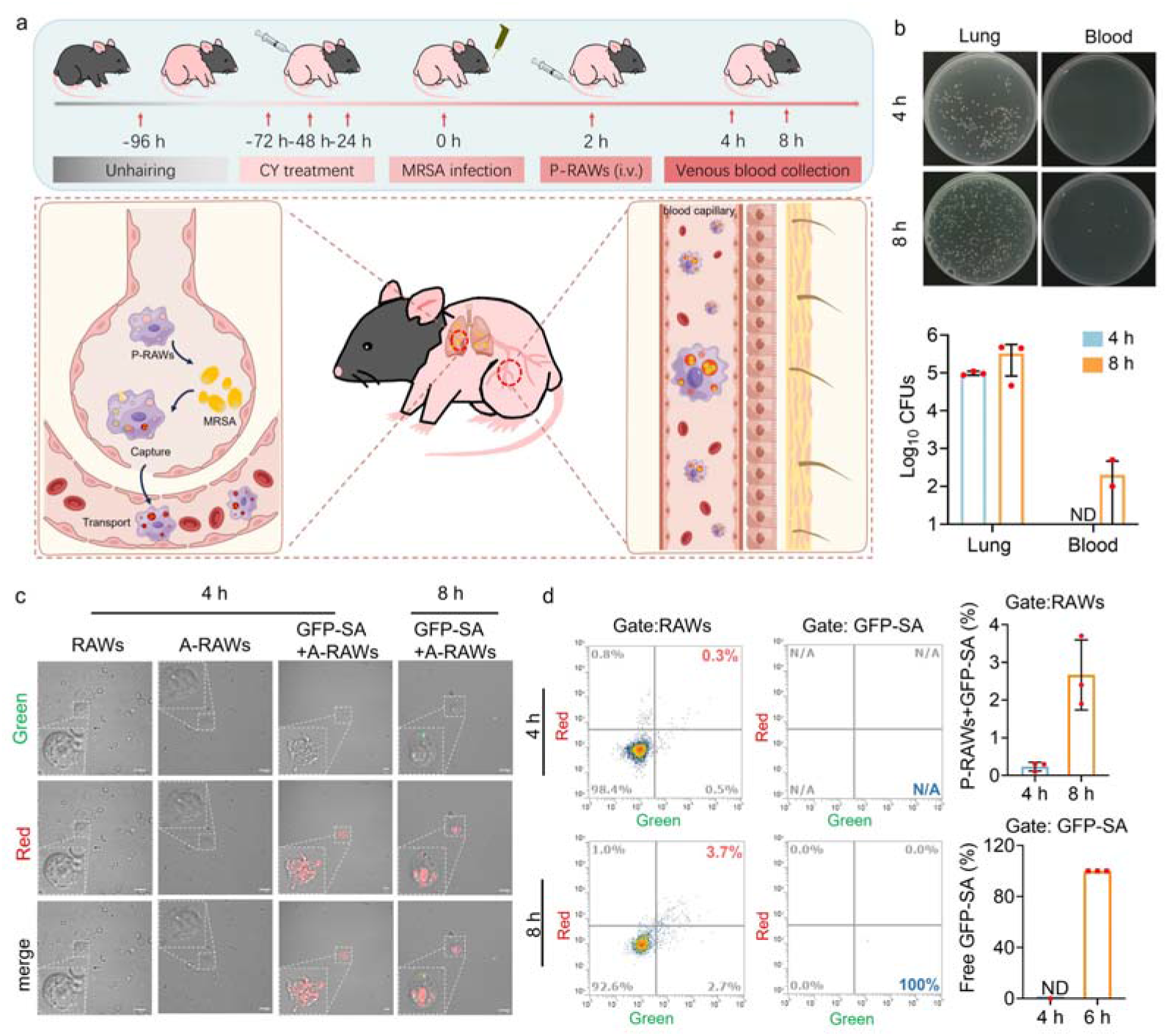
(a) Schematic diagram of live drug distant transport of GFP-SA in lung infections of immunodeficient mice. (b) Plate counting of GFP-SA load in the lung and blood at 4 h and 8 h after injection of GFP-SA. (c) Confocal imaging in blood at 4 h and 8 h after injection of GFP-SA to examine live drug transporting GFP-SA from the lungs. Bar: 10 µm. (d) Flow cytometry in blood at 4 h and 8 h after injection of GFP-SA to examine live drug transporting GFP-SA from the lungs.

We first induced an immunocompromised state of mice by treating healthy mice with cyclophosphamide (CY) for three consecutive days. To facilitate the monitoring through fluorescence imaging and flow cytometry, we used *Staphylococcus aureus* expressing green fluorescent protein (GFP-SA) to build the lung infection via intranasal administration (Figure 6a). For confocal imaging, we set up 4 groups, named RAWs (4 h), A-RAWs (4 h), GFP-SA+A-RAWs (4 h) and GFP-SA+A-RAWs (8 h). For the RAWs (4 h) group and A-RAWs group (4 h), we injected RAWs and A-RAWs into immunodeficient mice (without bacteria) via intravenous administration synchronously with the test group, respectively, and collected blood samples 2 h later for imaging. For two GFP-SA+A-RAWs groups, we first administrated GFP-SA into immunodeficient mice via intranasal administration. We then injected A-RAWs into the mice intravenously 2 h after the infection. We finally collected blood samples 2 h and 6 h after the administration of A-RAWs through the tail vein for imaging (Figure 6a). The blood samples were diluted with PBS (V_blood_: V_PBS_=1:1000) and imaged using a confocal microscope to identify A-RAWs and GFP-SA. As shown in Figure 6c, we observed neither green fluorescence from bacteria (not injected into mice) nor red fluorescence from Lyso710A (not injected into mice) in the RAWs group. For the A-RAWs group, we observed neither green fluorescence from bacteria (not injected into mice) nor red fluorescence from Lyso710A (although A-RAWs are injected into mice). We inferred that A-RAWs maintained in standby mode, namely, the fluorescence and photodynamic effect of Lyso710A is in the off-state, in the absence of bacteria. For the GFP-SA+A-RAWs (4 h) group, we observed no green fluorescence from free bacteria (though GFP-SA is administrated into mice), as no bacteria spread out of the lung at this stage of infection. However, we identified the presence of A-RAWs with bacteria intracellular as demonstrated by colocalization of the red fluorescence from Lyso710A in on-state and the green fluorescence from GFP-SA (Figure 6c, GFP-SA+A-RAWs group, 4 h). We inferred that these A-RAWs captured bacteria in the lungs and transported them to the blood system. For the GFP-SA+A-RAWs (8 h) group, we observed A-RAWs with bacteria intracellular as well as free bacteria. It may indicate the presence of A-RAWs that phagocytic GFP-SA in the blood. However, we could not tell when and where (in the lung or the blood) A-RAWs phagocytose GFP-SA (Fig 6c).

To obtain statistical results, we performed the above experiments using flow cytometry. The detailed setting of control groups is listed in the supporting information. We focus on measuring the population of A-RAWs with bacteria intracellular (double positive in the green and red channels) and free bacteria (single positive in the green channel) in the test group CFP-SA+A-RAWs. Due to the difference in size between RAWs and GFP-SA, separate gates for GFP-SA (Gate: GFP-SA) and RAWs (Gate: RAWs) were created and set up to independently identify A-RAWs and GFP-SA controls to determine bacterial and cellular signals. Note that the presence of free GFP-SA is not reflected by the green single positive signal in Gate: RAWs, but by the green single positive signal in Gate: GFP-SA. Generally, the flow cytometry results are consistent with the imaging results. As shown in Figure 6d, in the A-RAWs+GFP-SA 4 h group, the proportion of red fluorescent single positive signal was 98.77%, and the proportion of red and green fluorescent double positive signal was 0.23% (Fig 6d red frame). Moreover, no green fluorescent single positive signal was shown in the Free GFP-SA group at this time (Fig 6d blue frame). This indicates that there is a small amount of A-RAWs in the blood, which is activated after capturing the bacteria. They are likely to be transported in the blood circulation from the site of infection in the lungs. In the A-RAWs+GFP-SA 8 h group, the proportion of red fluorescent single positive signal was 92.17%, and the proportion of red and green fluorescent double positive signal was 2.67%. The number of double-positive cells was significantly higher than that at 4h. At this time, the Free GFP-SA group also showed the presence of a green fluorescent single positive signal, but the number was very small (Fig 6d). These results demonstrate the natural circulation and long-range bacterial trapping and killing capability of A-RAWs, effectively controlling bacterial infection.

## Conclusion

In this work, we propose a “live drug” that enables photodynamic therapy of systemic bacterial infection by bathing in the sunshine. To the best of our knowledge, this is the first report of photodynamic therapy of deep-tissue disease by irradiating the skin using sunlight. This strategy takes advantage of the therapeutic efficiency of photodynamic therapy and the innate chemotaxis of macrophages. On the one hand, the “off-on” mechanism of the photodynamic effect of Lyso710A makes our therapy highly bacteria-specific, and safe for healthy cells in the neighborhood. On the other hand, macrophages are armed with Lyso710A in their lysosomes, as the smart hunters for the bacteria. Our results demonstrate that this “live drug” enables an accurate delivery of bacteria-activable photosensitizer (Lyso710A) to the surface of bacteria, an efficient transportation of bacteria to the epidermis layer through the bloodstream, therefore an accurate elimination of bacteria in the sunshine, and restricts the photo-damage away from healthy tissue. Generally, the smart targeting ability of macrophages endows high efficiency, bio-safety and operability of this novel style of photodynamic therapy and leads to a cure of the systemic bacterial infection at 100% in immunodeficient mice model. We will focus on optimizing the dose of photosensitizer and quantifying the transportation efficiency of macrophages in our future work.

## Supporting Information

The Supporting Information is available free of charge at AUTHOR INFORMATION

## Corresponding Author

**Xinfu Zhang** − State Key Laboratory of Fine Chemicals, Frontiers Science Center for Smart Materials Oriented Chemical Engineering, Dalian University of Technology, Dalian 116024, China.

**Yi Xiao** − State Key Laboratory of Fine Chemicals, Frontiers Science Center for Smart Materials Oriented Chemical Engineering, Dalian University of Technology, Dalian 116024, China.

## Acknowledgments

This work was supported by the National Natural Science Foundation of China (No. 22278059, 22174009, 22078047 and 21901031), Fundamental Research Funds for the Central Universities (No. DUT22LAB601 and DUT22LAB608), Science and Technology Foundation of Liaoning Province (2020-YQ-08).

## Author contributions

Conceptualization of the study was done by X.Z.; the experimental design was carried out by Z.W., X.Z. and Y.X.; chemical synthesis was carried out by Z.W.; cell experiments were carried out by Z.W.; the mouse model was constructed by Z.W.; analysis of results was done by X.Z., Z.W., L.W., L.Z. and Y.X.; the manuscript was written by X.Z., Z.W. and Y.X.; the project was conceived and supervised by X.Z. and Y.X.

