## Supporting information for "Armed Macrophages as Hunters for Photodynamic Therapy of Systemic Bacterial Infections by Bathing in the Sunshine"

### 1. Materials and instruments.

All chemicals used in the experiment were purchased from commercial suppliers (Bidepharm or Energy Chemical) were not subjected to further purification. The following reagents were purchased from Yeasen Biotech Co., Ltd.: Hoechst 33342, LysoTracker® Green DND-26, 2′, 7′-dichlorodihydrofluorescein diacetate (DCFH-DA), and Calcein-AM/PI Double Stain Kit. 3-(4,5-dimethylthiazol-2-yl)-2,5-diphenyl-2h-tetrazoliubromide (MTT) was purchased from Bidepharm.

The $^{1}$H-NMR spectra were measured using TMS as an internal reference in $CDCl_3$ on a 400 or 500 MHz spectrometer. The $^{1}$H-NMR spectra were measured using TMS as an internal reference in $CDCl_3$ on a 400 or 500 MHz spectrometer. Additionally, the $^{13}$C-NMR spectrum was registered on a 100 MHz spectrometer. Mass spectra were obtained using an Agilent LC/Q-TOF MS.

UV-vis-NIR spectra were obtained using either the Agilent 8453 UV-visible or Lambda 750S spectroscopy system with a path length of 1 cm. Fluorescence images were acquired using an Olympus Fluoview FV1000 confocal laser scanning microscope. Fluorescence imaging of small animals was performed using the NightOWL II LB983 living imaging system.

### 2. Synthesis of Lyso710A.

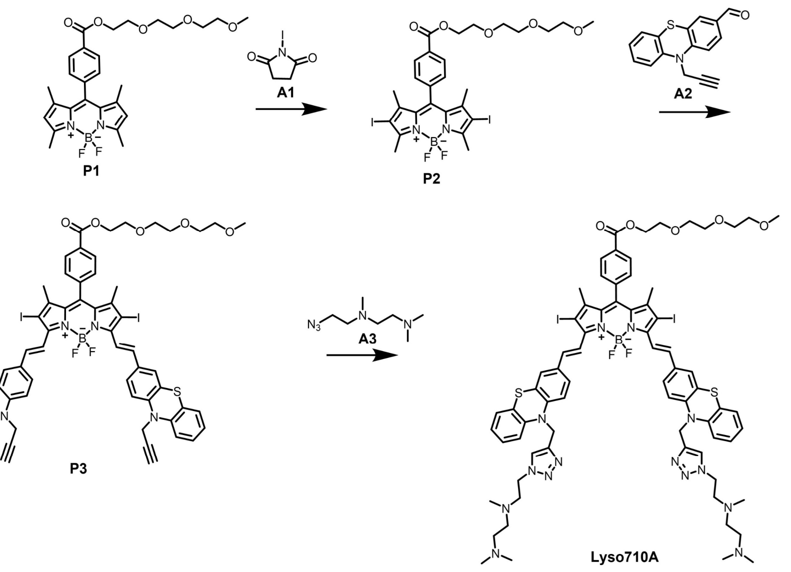


**SI Figure 1.** Synthetic route of Lyso710A.

P1, P2, A2, and A3 were synthesized according to literature methods[1, 2].

2.1 Synthesis of P3.

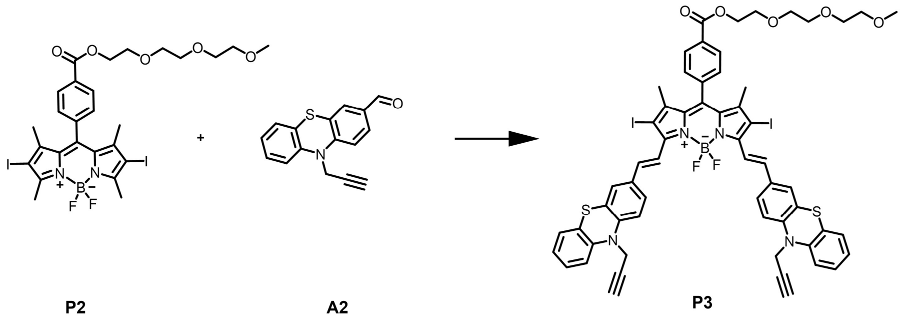


P2 (200 mg, 261.04 μmol) and A2 (173.15 mg, 652.60 μmol) were dissolved in 5 ml of anhydrous toluene. Then, 0.1 ml of piperidine and 0.1 ml of glacial acetic acid were added successively. The mixture was refluxed at boiling point and the reaction progress was monitored by TLC. After completion, the solution was diluted with deionized water and extracted with $CH_2Cl_2$ (3X). The combined organic solvent layer was dried with $MgSO_4$ and concentrated. The green solid P3 (190 mg, 150.70 μmol) obtained in a 57.73% yield after purification of the crude product through flash chromatography (silica gel). $^1$H NMR (400 MHz, $CDCl_3$) δ 8.23 (d, J = 8.2 Hz, 2H), 8.09 (d, J = 16.6 Hz, 2H), 7.60 (s, 1H), 7.56 (s, 1H), 7.52 (dd, J = 8.5, 2.0 Hz, 2H), 7.42 (t, J = 1.7 Hz, 3H), 7.41 (d, J = 2.5 Hz, 1H), 7.26 (s, 1H), 7.23 (s, 1H), 7.21 (d, J = 1.8 Hz, 3H), 7.20 (d, J = 1.4 Hz, 1H), 7.17 – 7.13 (m, 2H), 6.97 (ddd, J = 8.2, 6.4, 2.3 Hz, 2H), 4.55 (s, 4H), 4.53 (s, 2H), 3.92 – 3.87 (m,

2H), 3.77 – 3.74 (m, 2H), 3.73 – 3.70 (m, 2H), 3.69 – 3.66 (m, 2H), 3.58 – 3.55 (m, 2H), 3.39 (s, 3H), 2.49 (t, J = 2.1 Hz, 2H), 2.07 (s, 2H), 1.43 (s, 4H). HRMS (ESI, m/z): calcd for $C_{59}H_{49}BF_2I_2N_4O_5S_2$, 1260.1295; found $[M+Na]^+$, 1283.1191.

2.2 Synthesis of Lyso710A.

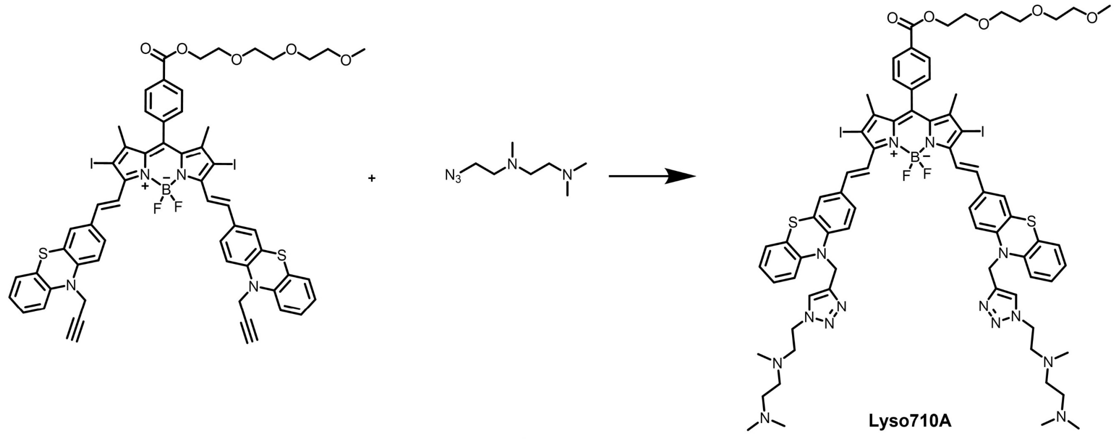


P3 A3

P3 (100 mg, 79.31 μmol), A3 (67.91 mg, 396.57 μmol), sodium ascorbate (3.14 mg, 15.86 μmol), and $CuSO_4$ $5H_2O$ (3.96 mg, 15.86 μmol) were added to a mixture of THF and $H_2O$ (v/v 3:1) under argon protection. The resulting mixture was stirred at room temperature for 24 hours. The solvent was then distilled off under reduced pressure, and the product was purified by silica gel column chromatography using $CH_2Cl_2$:EtOH (35:1, v/v) as the developing solvent. This yielded 97 mg (60.50 μmol) of dark green solid Lyso710A, with a 76.28% yield.$^1$H NMR (400 MHz, $CDCl_3$) δ 8.24 (d, J = 7.9 Hz, 2H), 7.99 (d, J = 16.4 Hz, 2H), 7.68 (s, 2H), 7.50 (s, 1H), 7.46 (s, 1H), 7.41 (d, J = 7.9 Hz, 2H), 7.36 (s, 2H), 7.32 (d, J = 8.8 Hz, 2H), 7.15 (d, J = 7.5 Hz, 2H), 7.10 (d, J = 7.7 Hz, 2H), 6.96 (t, J = 7.5 Hz, 2H), 6.86 (d, J = 14.7 Hz, 4H), 5.23 (s, 4H), 4.55 (s, 2H), 4.43 (t, J = 5.5 Hz, 4H), 3.90 (s, 2H), 3.76 (d, J = 5.1 Hz, 2H), 3.71 (d, J = 5.2 Hz, 2H), 3.69 – 3.65 (m, 2H), 3.57 (d, J = 4.8 Hz, 2H), 3.39 (s, 3H), 2.85 – 2.78 (m, 4H), 2.64 – 2.59 (m, 4H), 2.51 (s, 4H), 2.34 (s, 12H), 2.14 (s, 6H), 1.43 (s, 6H). HRMS (ESI, m/z): calcd for $C_{73}H_{85}BF_2I_2N_{14}O_5S_2$, 1602.4263; found $[M+2H]^{2+}$, 802.2205.

### 3. Optical responses of Lyso710A

Lyso710A was diluted to 5 μM in a 0.1% Triton X-100/PBS solution. The UV-vis and fluorescence spectra were recorded after adding analytes to the solutions. The spectroscopic experiments were conducted at room temperature.

### 4. In vitro oxygen ($^1O_2$) detection of Lyso710A.

The singlet oxygen capacity of Lyso710A was assessed using 1, 3- diphenylisobenzofuran (DPBF). The capacity of Lyso710A to produce singlet oxygen was evaluated using 1,3-diphenylisobenzofuran (DPBF). The absorbance of DPBF at 415 nm was adjusted to approximately 1.0 in a 0.1% Triton X-100/PBS solution with or without 0.1% trifluoroacetic acid (TFA). The cuvette was exposed to 660 nm monochromatic light for different durations, and absorption spectra were measured promptly. The absorbance slopes of DPBF at 415 nm were measured against irradiation time to compare the generation ability of $^1O_2$ with and without acid.

$^1O_2$ quantum yield determination of Lyso710A (ps) using Methylene Blue (MB) in DCM as reference. $\Phi_\Delta$ was calculated by the following equation:

$$\Phi_{\Delta(ps)} = \Phi_{(MB)}(\frac{A_{ps}}{A_{MB}})(\frac{F_{MB}}{F_{ps}})(\frac{PF_{MB}}{PF_{ps}})$$

Where A is the slope of a plot of the change in absorption of DPBF at 415 nm, F is the absorption correction factor, which can be determined by $F = 1 - 10^{-OD}$, and PF is an absorbed photonic flux (μ Einstein $dm^{-3}$ $s^{-1}$). $\Phi_{MB}$ (0.57) is the $^1O_2$ quantum yield of MB in DCM.

### 5. Preparation of stock solutions or generation of ROS

(1) $ClO^-$

The NaClO solution was appropriately diluted in 0.1 M NaOH solution. The concentration of $ClO^-$ was determined using the molar extinction coefficient at 292 nm (350 $M^{-1}$ $cm^{-1}$). Subsequently, the $ClO^-$ stock solution was prepared.

(2) $NO_2^-$

Ultrapure deionized water (25 mL) was used to dissolve 0.8625g of sodium nitrite, resulting in the concentration of $NO_2^-$.

(3) $TBO^•$

The Fenton reaction of 50 mM TBHP with 50 mM $FeSO_4$ solution generated the tert-butoxy radical ($TBO^•$).

(4) $NO^•$

Nitric oxide was produced from Sodium Nitroferricyanide (III) Dihydrate (SNP). To generate $NO^•$,

14.8975mg of SNP was added to 10mL of Ultrapure deionized water and stirred for 30 minutes at 25°C, resulting in a concentration of 5mM.

(5) $^1O_2$

At room temperature, sodium phosphate buffer at pH 7.4 was used to mix $H_2O_2$ and $ClO^-$ at the same molar concentrations.

(6) TBHP

Ultrapure deionized water (40 mL) was used to deliver *Tert*-butyl hydroperoxide (TBHP) at a concentration of 50 mM.

(7) $H_2O_2$

The $H_2O_2$ was appropriately diluted in Ultrapure deionized water, and its concentration was determined using the molar extinction coefficient at 240 nm (43.6 $M^{-1}$ $cm^{-1}$). Subsequently, the $H_2O_2$ stock solution was prepared.

(8) $^{\bullet}OH$

The $H_2O_2$ solution was added to a sodium phosphate buffer with a pH of 7.4, followed by the addition of a $FeSO_4$ solution (50 μM) at room temperature. This resulted in the generation of $^{\bullet}OH$ through the Fenton reaction between $Fe^{2+}$ and $H_2O_2$.

### 6. Cell and Culture Conditions.

Raw264.7 cells were obtained from the Institute of Basic Sciences (IBMS) of the Chinese Academy of Medical Sciences (CAMS). The cells were cultured in DMEM supplemented with 10% fetal calf serum (FBS) and 0.1 mg $ml^{-1}$ streptomycin at 37°C in a 95% air and 5% $CO_2$ environment. For imaging purposes, all cell types were cultured on 35 mm glass-bottom culture dishes for 12-24 hours.

### 7. The standard method to prepare A-RAWs

Raw264.7 cells in the exponential growth phase were cultured on 35 mm confocal dishes at a density of $5\times10^4$ cells per dish for 24 hours. Subsequently, the cells were incubated with 5 μM Lyso710A for 3 hours at 37 °C under 5% $CO_2$. After incubation, the cells were washed three times with PBS. A-RAWs were obtained by trypsin digestion.

### 8. Confocal Fluorescence Imaging.

Raw264.7 cells in the exponential growth phase were cultured on 35 mm glass-bottom culture dishes until they reached 80% confluency, which took 24-36 hours. The cells were then washed three times with PBS and incubated with 1 mL DMEM containing Lyso710A (5 μM) at 37°C in an atmosphere of 95% air with 5% $CO_2$ for 3 hours. A-RAWs were activated using HClO. The cells were further stained with DND-26 (1 μM) and visualized using laser confocal microscopy. Lyso710A was excited at a wavelength of 635 nm, while DND-26 was excited at 488 nm. The emission wavelength range was 655-755 nm for Lyso710A and 510-550 nm for DND-26. The emission wavelength range was 655-755 nm for Lyso710A and 510-550 nm for DND-26.

**9. Intracellular ROS Detection.**

Raw264.7 cells were plated onto 35 mm confocal dishes and incubated with 5 μM Lyso710A for 3 hours. Subsequently, they were stained with 1 μM DCFH-DA for an additional 30 minutes. Following irradiation for 15 minutes with LED daylight (70 mW/cm$^2$), the fluorescence channels were collected, and their average intensity was calculated.

**10. Confocal imaging of intracellular phototoxicity.**

Raw264.7 cells were plated onto 35 mm confocal dishes and incubated with 5 μM Lyso710A for 3 h. After irradiating for 15 min with LED daylight (70 mW/cm$^2$), cells were stained with Calcein (AM)/propidium iodide (PI) Apoptosis Detection Kit according to the manufacturer's instructions. The cell apoptosis was visualized by fluorescence microscopy with an excitation wavelength of 488 nm and 559 nm.

**11. MTT assay.**

Concentration-dependent cell viability.

Raw264.7 cells were seeded onto 96-well plates at a density of $3\times10^3$ cells per well and cultured for 24 hours. Lyso710A was added at varying concentrations and incubated for 3 hours. The cells were then washed three times with PBS. For dark toxicity assessment experiments, the 96-well plates were incubated for 24 hours in the absence of light. For phototoxicity assessment, the 96-well plates were irradiated with LED daylight (70 mW/cm$^2$) for 15 minutes. After irradiation, the cells were incubated for an additional 24 hours. Cell viability was measured using the MTT assay.

### 12. Safety evaluation of A-RAWs antimicrobial strategy in simulated normal tissues.

Raw264.7 cells were seeded in 35 mm glass-bottom culture dishes and cultured for 24 hours. They were then incubated with 5 μM Lyso710A for 3 hours at 37 °C under 5% $CO_2$. The cells were subsequently washed three times with PBS. COS-7 cells were seeded in the same 35 mm glass-bottom culture dishes and cultured for 24 hours. After being irradiated for 10, 20, or 30 minutes with LED daylight (70 $mW/cm^2$), the cells were stained with the Calcein (AM)/propidium iodide (PI) Apoptosis Detection Kit according to the manufacturer's instructions. Cell apoptosis was visualized using fluorescence microscopy with an excitation wavelength of 488 nm.

### 13. Bacterial culture.

The Methicillin-resistant *Staphylococcus aureus* (MRSA) strain, ATCC 43300, was obtained from American-type cultures (ATCC, Manassas, VA). The bacteria were cultured overnight at 37°C in 20 mL of LB medium (containing Tryptone 10 g/L, Yeast extract 5 g/L, and NaCl 10 g/L). Once the bacteria reach the logarithmic growth phase, rinse them three times with phosphate-buffered saline (PBS). Then, stain them with Mem-SQAC (1 μM) for 30 minutes. Rinse them again with PBS three times and finally dilute them with DMEM or MEM medium to achieve the desired bacterial concentration for the multiplicity of infections.

### 14. Survival of bacteria within A-RAWs post phototherapy.

The MRSA was diluted and added to A-RAWs thin monolayer culture. After 30 min, the cells were treated with 100 μg/mL gentamicin sulfate (Sigma-Aldrich, G1914) for 1 h to kill extracellular bacteria. All groups were separately covered under 1, 2 mm chicken tissue irradiated upon LED daylight (70 $mW/cm^2$) for 40 min. After incubation, the cells were washed with PBS and further incubated in 10% FCS-DMEM. Finally, wash the cells with PBS to wash away extracellular bacteria and dead cells, add 0.5% Triton X-100 (Sigma-Aldrich, X100) to lyse, and apply a series of dilutions on LB agar (Sigma-Aldrich, 22092 and BD Biosciences, 214010) for bacterial counts. Specifically, 10 ul of blood and homogenates of major organs were collected from mice and diluted $10^1$, $10^2$, $10^3$, $10^{4,}$ and $10^5$ times respectively. Plates were coated with the original solution and each dilution. The results of plate counting after dilution followed the dilution pattern.

**15. Construction of systemic bacterial infections model in immunodeficiency mice.**

Female C57 BL/6 mice aged 6-7 weeks were purchased from Dalian Medical University's Shanghai Experimental Animal Center. The animal experiments conducted in this study were approved by the Animal Care and Use Committee of Dalian Medical University. Intraperitoneal administration of cyclophosphamide was used to construct mouse immunodeficiency models, a widely studied and established method. C57BL/6 mice were intraperitoneally injected with 100 mg/Kg Cyclophosphamide (CY) for bacterial infection over three consecutive days. The development of immune deficiency and health recovery in mice was assessed by monitoring mouse WBCs and LYMs through blood routine tests.

For the construction of A-RAWs: Raw264.7 cells were seeded in a T75 cell culture flask at a density of $1\times10^8$ cells per flask and incubated for 12 hours. After that, the cells were stained with 5 μM Lyso710A in fresh culture medium for an additional 3 hours. The culture medium containing Lyso710A was then removed, and the cells were washed three times with PBS. A-RAWs were obtained by trypsin digestion and centrifugation. The cells were then dispersed in ethanol and left at room temperature for 20 minutes. The mixture was subsequently centrifuged at 800 g for 10 minutes to collect the supernatant. The amount of Lyso710A dye taken by cells was calculated as follows: Firstly, the absorption intensity of Lyso710A in the supernatant was measured using a UV-vis spectrophotometer (SI Figure 2a). Subsequently, the absorption intensity was converted to the corresponding concentration using a standard curve of the absorption intensity versus the concentration of Lyso710A in ethanol (SI Figure 2b). Lastly, the concentration of Lyso710A in $1\times10^8$ cells was calculated as 0.88 μM. We were able to obtain a mole amount of Lyso710A of $1.76\times10^{-9}$ mole in $1\times10^8$ cells. These A-RAWs in PBS were injected intravenously in each mouse infected with an equivalent dose of 0.14 mg/Kg.

For the construction of the mouse systemic bacterial infections model: mice were anesthetized with 10% chloral hydrate intraperitoneally. All mice were depilated all over their bodies. 50 μl of stained MRSA suspension ($1\times10^8$ CFU) was inoculated into the mouse blood system by tail vein injection.

For treatment procedure and data collection: At 2 h after MRSA injection, $1\times10^8$ A-RAWs were injected intravenously into each mouse. The whole body of mice was irradiated with sunlight or simulated sunlight (120 mW/cm$^2$, 6 hours/day) for three days after the bacteria injection, respectively. The mice's blood and main organs (heart, liver, spleen, lungs, and kidneys) were immediately collected, and bacterial CFU was quantified after three days of treatment. The survival and body weight (BWs) of mice was assessed every 48 h. After 30 days, the mice were sacrificed. The main organs were stained with H&E tissue to evaluate treatment outcomes.

**16. Construction of lung infections model in immunodeficiency mice.**

The operation steps for constructing a mouse immunodeficiency model are the same as those described above.

For the construction of the mouse lung bacterial infections model: mice were anesthetized with 10% chloral hydrate intraperitoneally. All mice were depilated all over their bodies. 50 μl of stained GFP-SA suspension ($1\times10^6$ CFU) was inoculated into the mouse lung by nasal feeding.

For confocal imaging of live drugs in the bloodstream: Control groups were set up by injecting RAWs and A-RAWs in the upper tail vein of immunodeficient mice, respectively. The experimental group was set up with $1 \times 10^6$ A-RAWs injected intravenously in each mouse after nasal administration of GFP-SA for 2 h on immunodeficient mice. Tail vein blood sampling (100ul) was performed on mice at 4 h, and 8 h after GFP-SA injection. Place in EDTA anticoagulation tube. Lysed red using erythrocyte lysate (Erythrocyte Lysate from Solarbio). Acquired cells were finally placed in an equal amount of PBS for confocal imaging. Place in EDTA anticoagulation tube. Lysed red using erythrocyte lysate. Acquired cells were finally placed in PBS for confocal imaging. The excitation wavelength for Lyso710A was 635 nm, while the excitation wavelength for GFP-SA was 488 nm. The emission wavelength was collected from 655 to 755 nm for Lyso710A and 510 to 550 nm for GFP-SA.

For flow cytometry of live drug in the bloodstream: to accurately assign A-RAWs and free GFP-SA populations and fluorescence signals in the blood of immunodeficient mice. We set up 4 control groups, RAWs and A-RAWs in vitro, only nasogastric fed GFP-SA in immunodeficient mice and injected with RAWs and A-RAWs by tail vein in immunodeficient mice without pneumonia, respectively, named RAWs, A-RAWs, CY+GFP-SA. CY+RAWs and CY+A-RAWs. In the test group, A-RAWs were injected into the tail vein of immunodeficient mice infected with pneumonia, named CY+A-RAWs+GFP-SA. Here, two hours after the infection (early systemic infection), mice were injected with RAWs or A-RAWs intravenously in the test group. Blood samples were collected through the tail vein at the 4th and 8th hour after the infection of bacteria for the test group. The blood samples were diluted with PBS (v:v=1:1000).

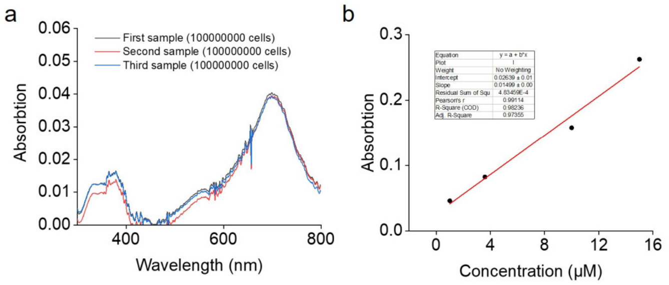


**SI Figure 2.** (a) Absorption intensity of Lyso710A in RAW264.7 (1×10$^8$ cells). (b) Standard curve of Absorption intensity fitting of Lyso710A in ethanol at different concentrations.

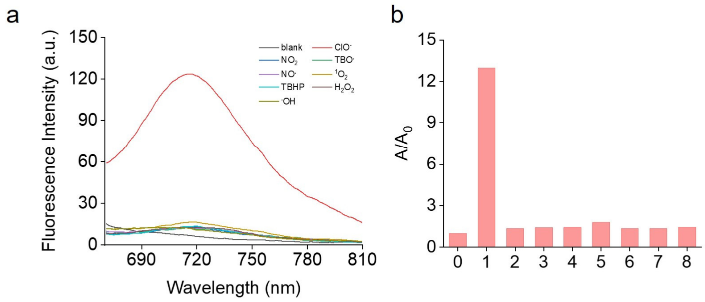


**SI Figure 3.** (a) Fluorescence intensity of Lyso710A (10 μM) upon treatment with ROS/RNS (1 mM) in 0.1% Triton X-100/PBS solution. (b) Fluorescence intensity ratio of Lyso710A (10 μM) upon treatment with ROS/RNS (1 mM) 0.1% Triton X-100/PBS solution.

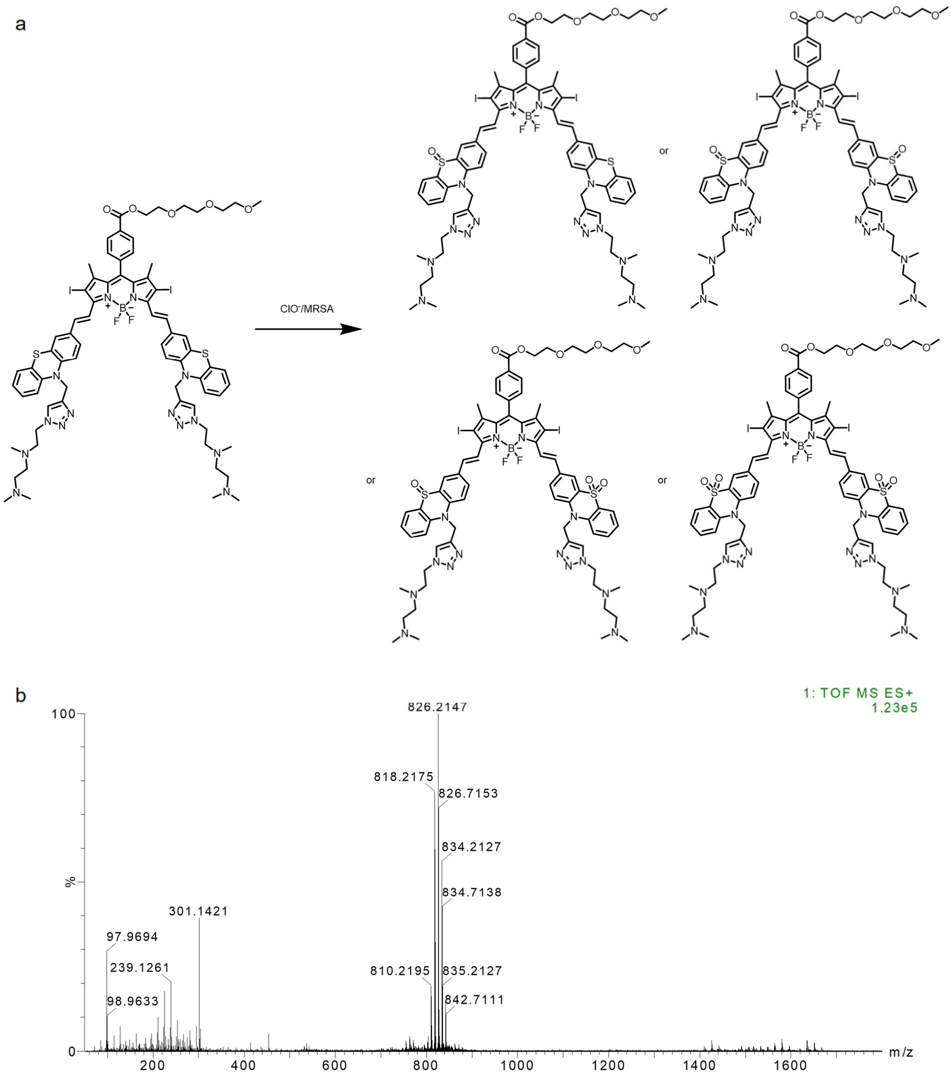


**SI Figure 4.** (a) Structural formula of Lyso710A oxidation product. (b) HRMS analysis of Lyso710A oxidation products induced by MRSA infestation in RAWs.

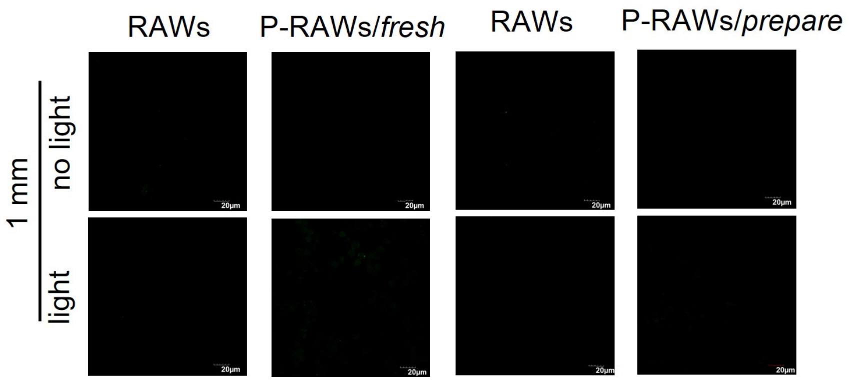


**SI Figure 5.** Fluorescence imaging of intracellular ROS production monitored by DCF-DA after light/dark incubation with A-RAWs/*fresh* and A-RAWs/*prepared* through 1 mm thickness of chicken meat, respectively. Bar: 20 µm.

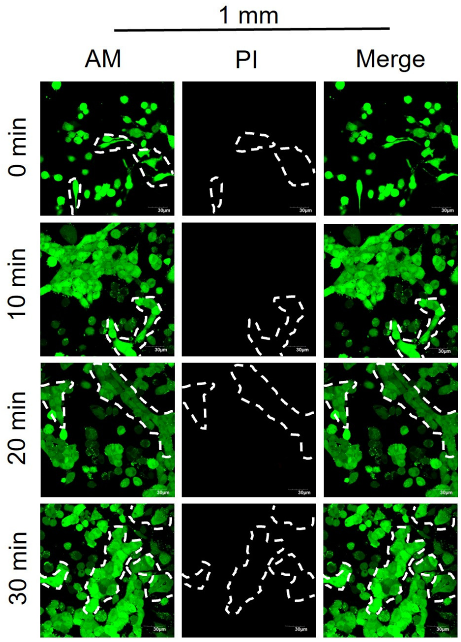


**SI Figure 6.** Calcein-AM/PI fluorescence imaging assesses the cell viability of co-cultured inactivated A-RAWs and COS-7 cells under simulated sunlight (70 mW/cm2, 10, 20, 30 min) through 1 mm thickness of chicken meat, respectively. The viability of cells was indicated through a Calcein-AM/PI co-staining assay. Bar: 30 μm.

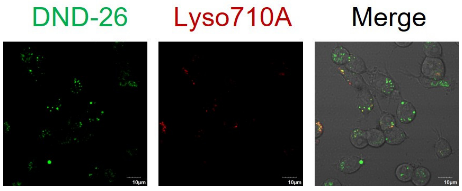


**SI Figure 7.** Fluorescence imaging of RAWs costained with Lyso710A and DND-26. Bar: 10 μm.

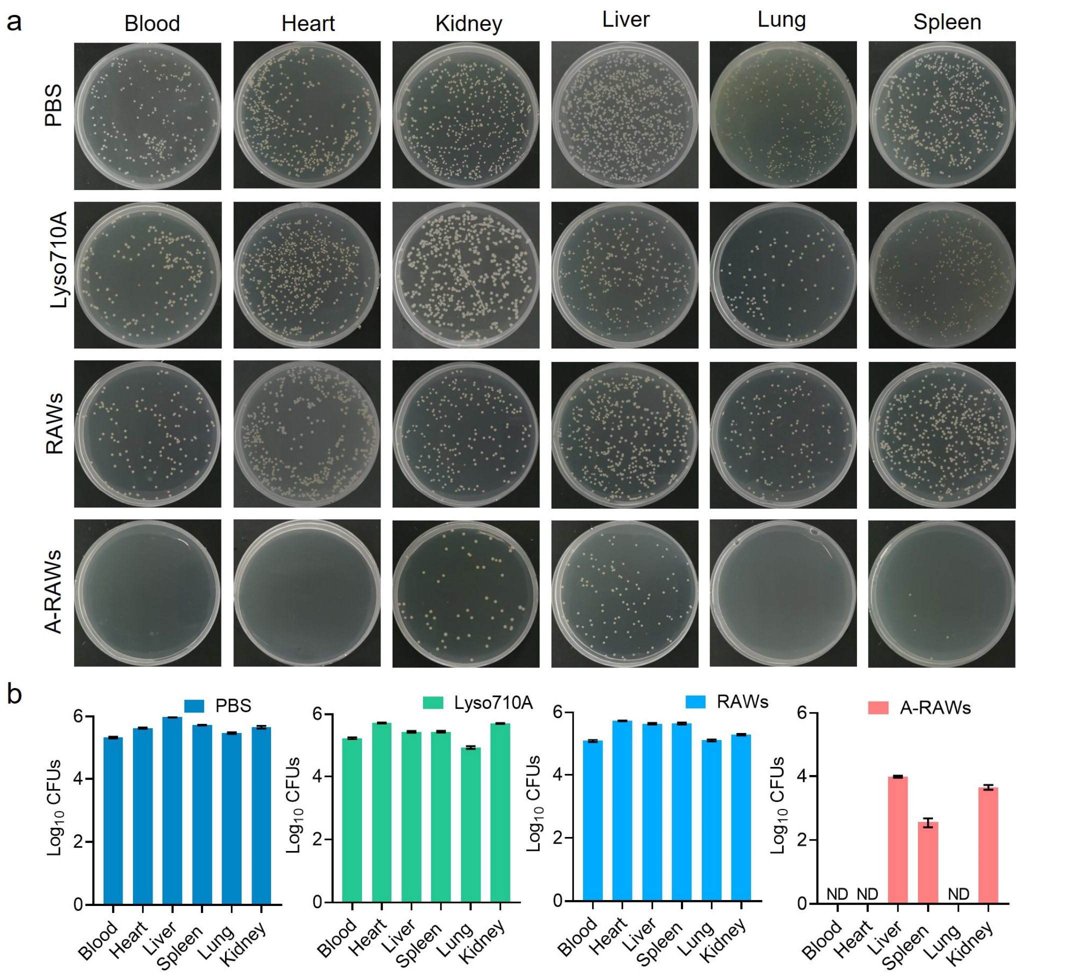


**SI Figure 8.** (a) Plate counting to assess bacterial load in major organs and blood in a systemic bacterial infection model in immunodeficient mice. (b) Bacterial load statistically in major organs and blood in an immunodeficient mice systemic bacterial infection model.

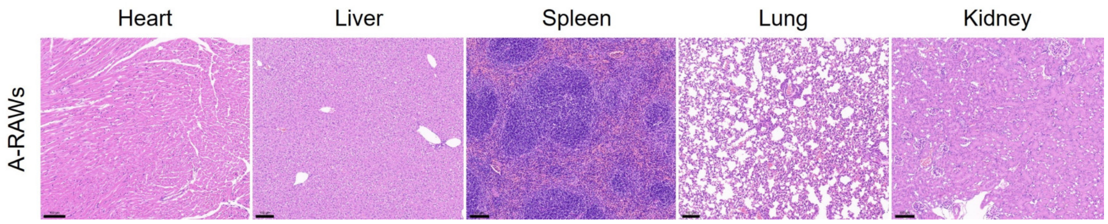


**SI Figure 9.** H&E staining of major organs in each group after 30 days in an immunodeficient mice systemic bacterial infection model. Bar: 100 nm.

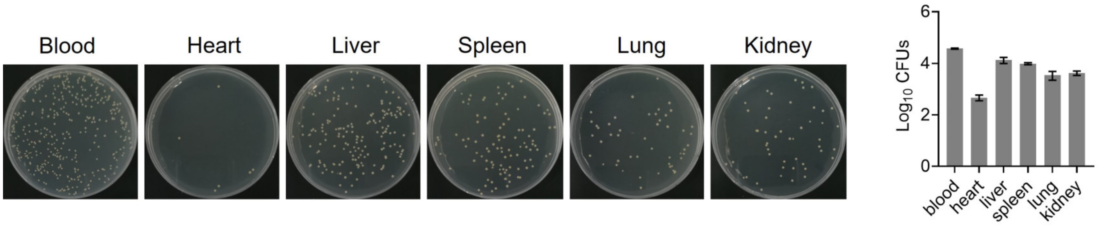


**SI Figure 10.** Plate counting to assess the bacterial load after 2 h of MRSA injection into the tail vein in a systemic bacterial infection model in immunodeficient mice.

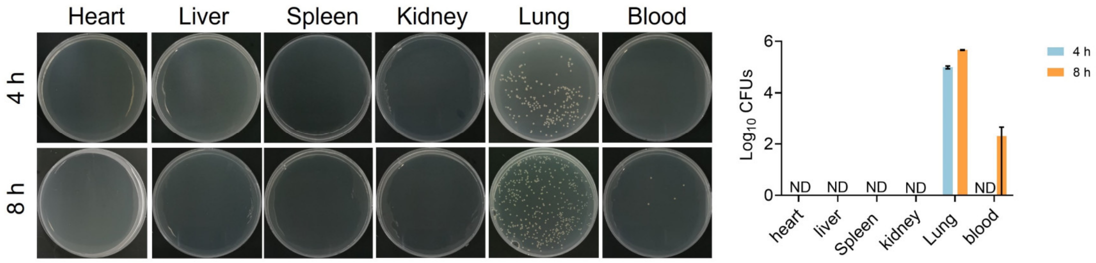


**SI Figure 11.** Plate counting to assess the bacterial load in blood at different moments without sunbathing in lung infections model of immunodeficient mice.

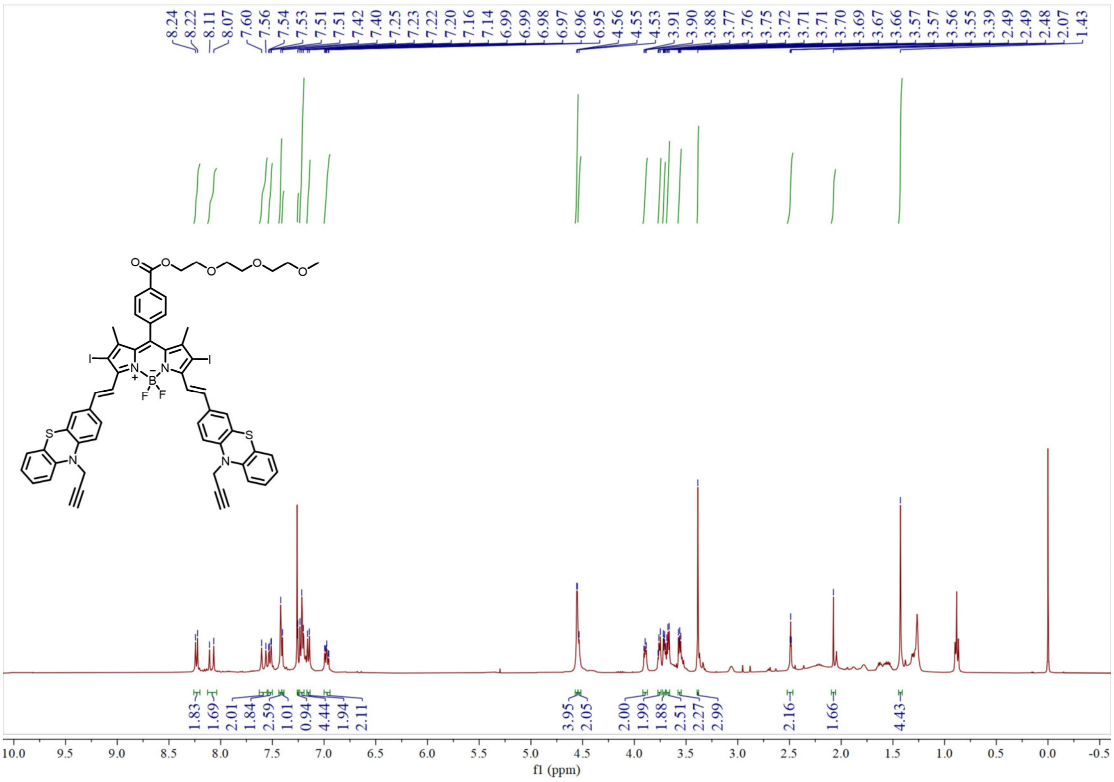


**SI Figure 12.** [1]H-NMR spectrum of P3 $CDCl_3$.

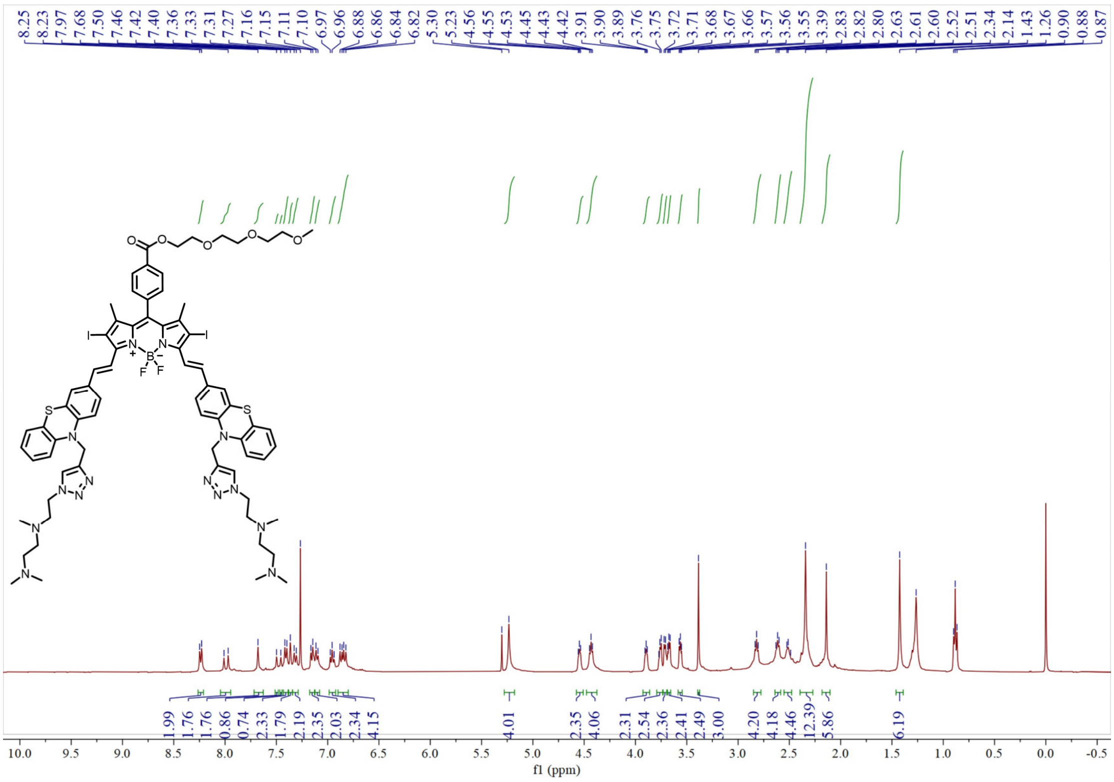

**SI Figure 13.** $^{1}$H-NMR spectrum of Lyso710A $CDCl_3$.

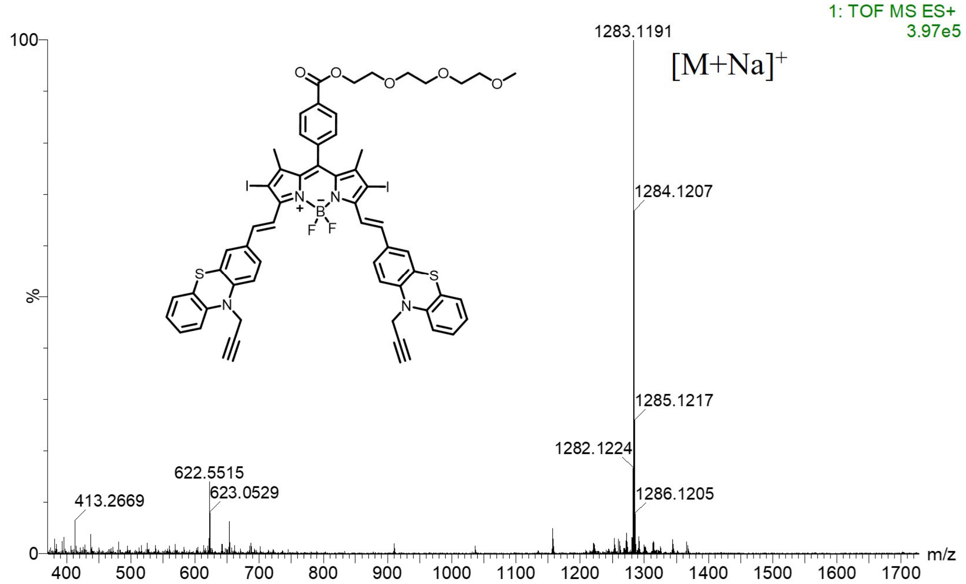


**SI Figure 14.** HRMS spectrum of P3.

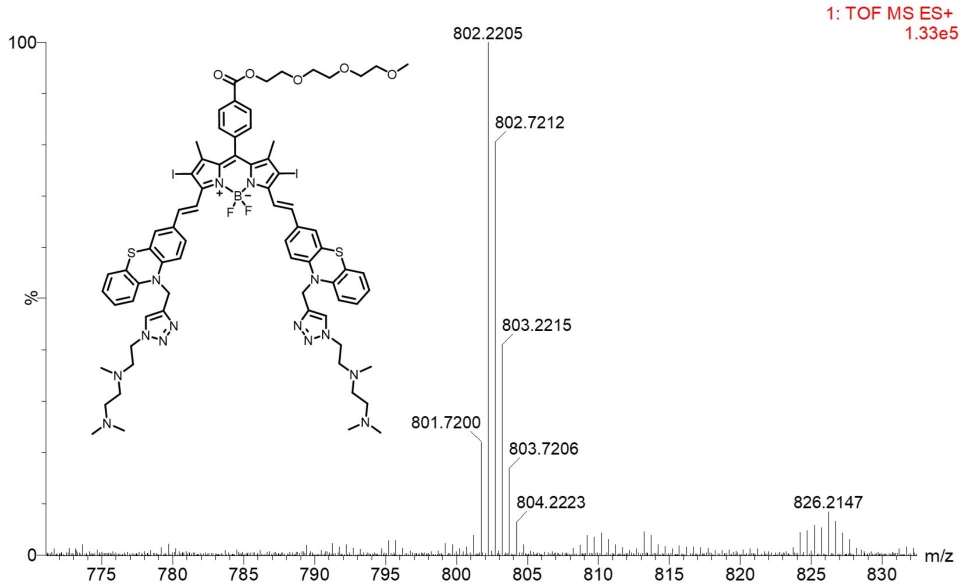


**SI Figure 15.** HRMS spectrum of Lyso710A.
